# The Evolution of Social Organization: Climate Influences Variation in Drosophilid Social Networks

**DOI:** 10.1101/776708

**Authors:** Jacob A. Jezovit, Rebecca Rooke, Jonathan Schneider, Joel D. Levine

## Abstract

Cultural norms, collective decisions, reproductive behaviour, and pathogen transmission all emerge from interaction patterns within animal social groups. These patterns of interaction support group-level phenomena that can influence an individual’s fitness. The aim of this study is to understand the evolution of social organization in Drosophila. Using a comparative ecological, phylogenetic and behavioural approach, we studied the different properties of social interaction networks (SINs) formed by 20 drosophilids and the different ways these species interact. We investigate whether animal network structures arise from common ancestry, a response to the species’ past ecological environment, other social behaviours, or a combination of these factors. We demonstrate that differences in past climate predicted the species’ current SIN properties. The drosophilid phylogeny offered no value to predicting species’ differences in SINs through phylogenetic signal tests. This suggests group-level social behaviours in drosophilid species are shaped by divergent climates. However, we find that the distance at which flies interact correlated with the drosophilid phylogeny, indicating that behavioural elements that comprise SINs have remained largely unchanged in their recent evolutionary history. We find a significant correlation of leg length to social distance, outlining the interdependence of anatomy and complex social structures. Although SINs display a complex evolutionary relationship across drosophilids, this study provides evidence of selective pressures acting on social behaviour in Drosophila. We speculate that conserved molecular mechanisms may be shared across drosophilids deep in their evolutionary history, similar to other pervasive mechanisms, like biological clocks.

## Introduction

Virtually all animals communicate and assemble into structured groups through a process called social organization [1]. Cooperating in these groups helps many animals overcome environmental stress and increase their fitness. For example, animals may avoid predation, increase foraging efficiency, access mates and locate optimal migration routes by collectively moving as a herd [2]. Understanding the nuances of social organization across animals has been enhanced by the application of social network analysis, since the structure of the network can predict and provide insight into group-level behaviours. For example, meerkat grooming networks can predict the likelihood of tuberculosis infection [3], and a finch’s position in their social network can influence how attractive they are to potential mates [4]. In *Drosophila melanogaster*, researchers have demonstrated that where a female lays her eggs is influenced by the structure of her social network [5]. Taken together, animal social networks can impact the health and behaviour of individual group members. Studying social networks can, therefore, provide a valuable tool for understanding the dynamics of social organization.

Despite its common label as a solitary insect, *D. melanogaster* displays a variety of social and collective behaviours. These include coordinating egg-laying sites [6, 7], trans-generational mate preference [8], collective feeding strategies [9], communicating the presence of predators/parasites to group members [10, 11] and collective escape responses [12]. Additionally, groups of *D. melanogaster* socially interact in a group setting that is structured and non-random. This was demonstrated through the previously published Social Interaction Network (SIN) assay [13]. A major finding from this assay was the intraspecific variation of a SIN property called betweeness centrality, a measurement of social network cohesiveness [14]. Betweeness centrality has also been reported to be heritable in humans [15], suggesting that the SIN assay may capture group-level phenotypes that are heritable in *Drosophila* and other organisms. With the SIN assay, we can measure distributions of replicated social networks, providing an excellent tool to study the sociality of *Drosophila*.

The *Drosophila* genus of insects is widespread throughout the world where groups of species have radiated into arid, temperate and tropical environments [16, 17]. The type of environment in which animals reside can correspond to differences in food availability, predation and species richness, among other things. These ecological variables have been shown to influence social network structure. For example, food abundance and predation pressure influence the interconnectedness of social networks in whales and guppies [18, 19]. Additionally, the taxonomic and phylogenetic relationships across drosophilids are well-established [20, 21]. One group measured thermal, cold and desiccation stress across 95 drosophilid species and found a strong phylogenetic correlation between cold tolerance and desiccation tolerance [22, 23]. Thus, this genus is well-suited for studying trait evolution in a macroevolutionary context.

Here, our aim is to apply comparative ecological, phylogenetic and behavioural approaches to understand the evolution of social organization in *Drosophila*. We hypothesized that drosophilid species vary in their social organization, measured through SINs, and that the species’ variation in SINs can result from three categorical influences (Figure 1): *behavioural elements of the network* (movement, social spacing, pairwise interactions; see Table 1), *phylogenetic signal* [24, 25], and *climatic selective pressures* [26]. In this report, we demonstrate how a combination of these three categories predicts drosophilid species’ variation in social behaviour. We reason that a broader comparative study of SINs across ecologically diverse drosophilid species enables us to investigate which factors have shaped social organization over macroevolutionary time. We show that the ability to form social networks in *Drosophila* is conserved and that their past environment predicts their present social structure. We suggest that the evolution of group-level interaction patterns may be supported by conserved mechanisms analogous to other pervasive mechanisms, such as biological clocks.

**Figure 1:**
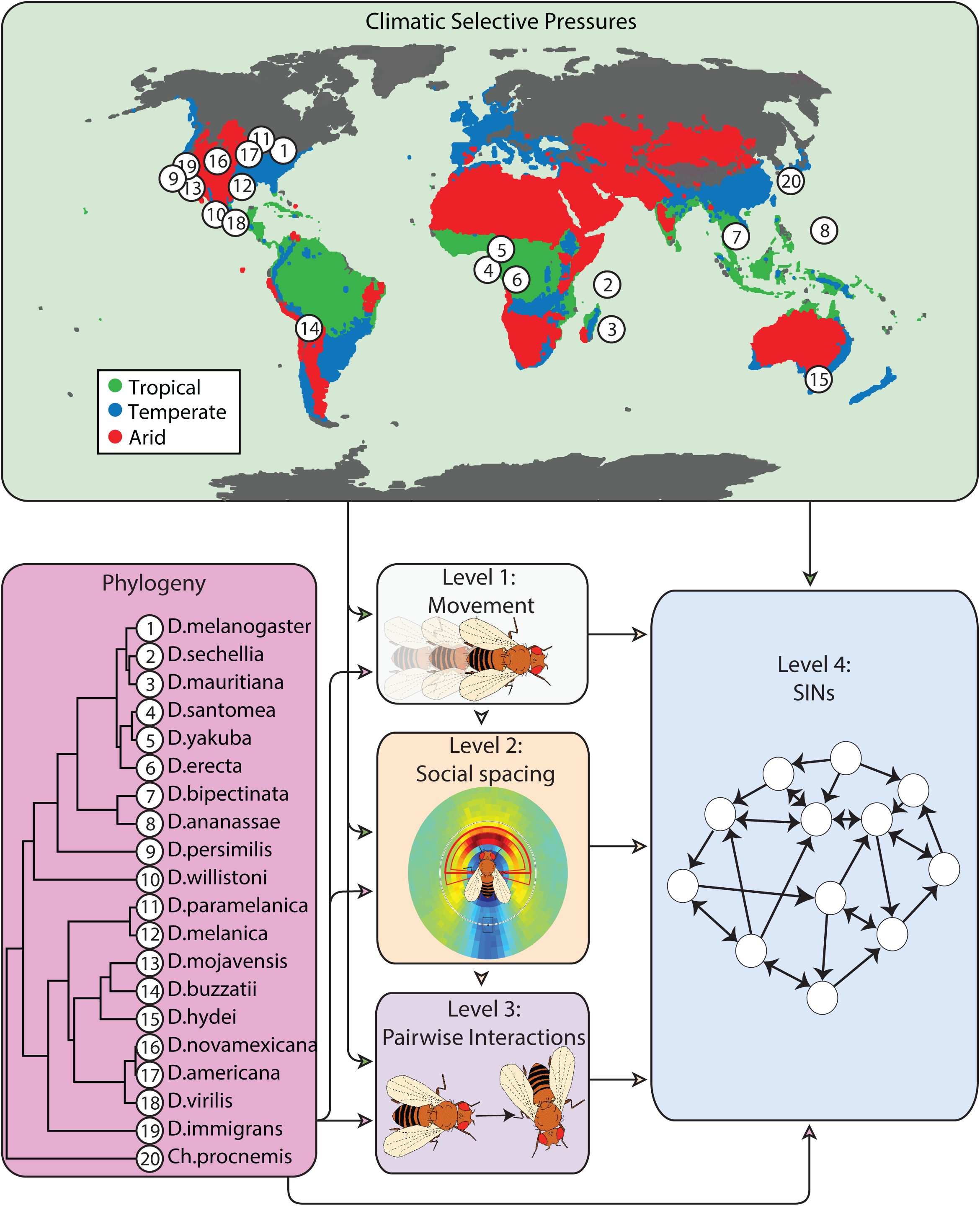
Hypothetical sources of influence on variation in drosophilid group level behaviours. Variation in Social Interaction Networks (SINs), may be influenced by past environmental selective pressures, differential common ancestry (phylogeny) and through a hierarchy of behavioural elements in order of increasing complexity: movement (level 1), the social spacing between interacting flies (level 2), and pairwise social interactions (level 3). See Table 1 for definitions of each of these elements. Arrows show how phylogeny, environment and behaviour are connected and indicate the relationships of phylogenetic and environmental influences on each behavioural variable. The phylogeny [21] lists the evolutionary history of the 20 drosophilid species we studied. The world map outlines the coordinates of the geographic origin of each species stock we studied. The colouration on the map indicate tropical regions (green), arid regions (red), and temperate regions (blue), based on the Koppen-Geiger climate classification [49]. See Table 1 for definitions of each behavioural measure within these three categories.

**Table 1:**
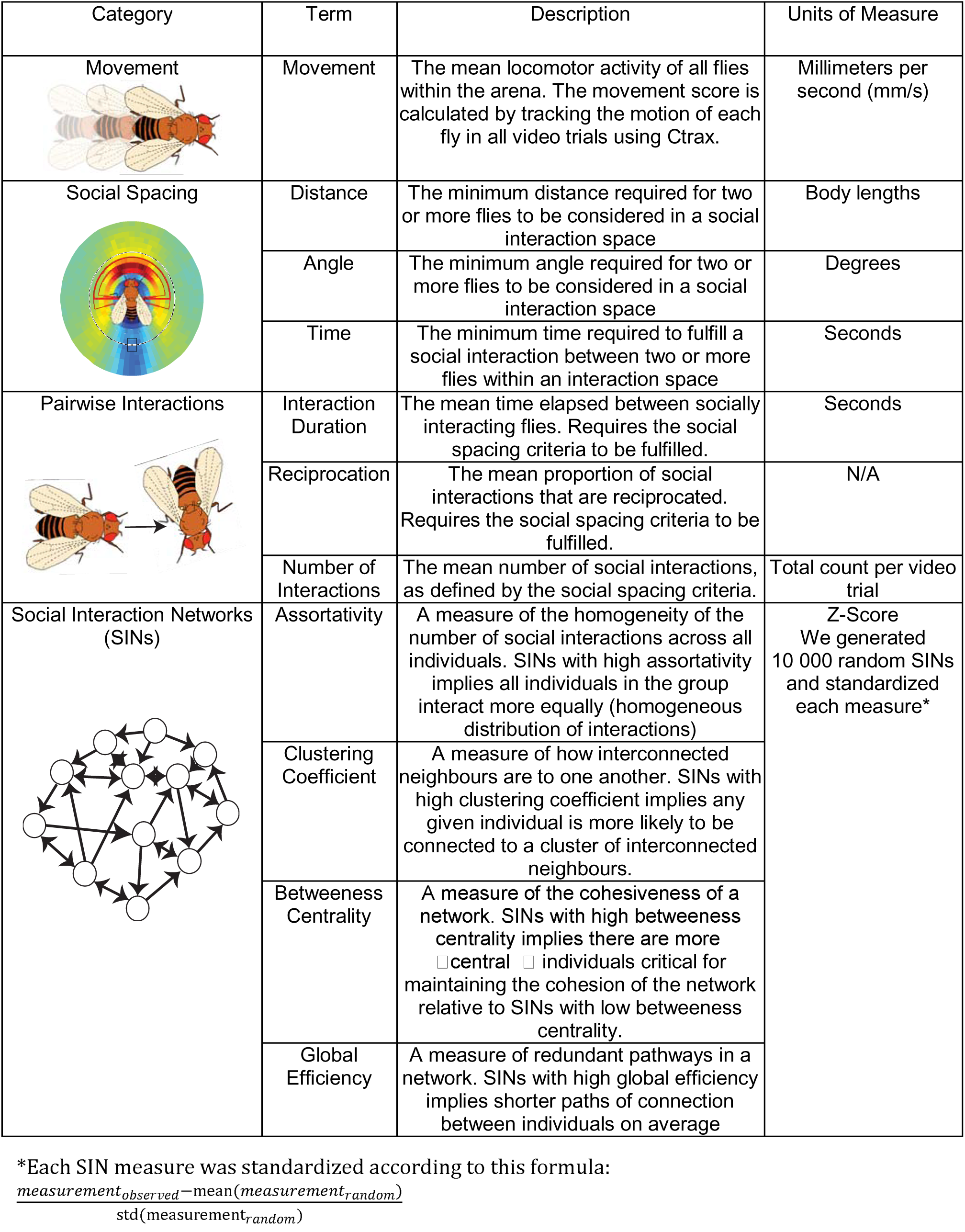
A glossary of all behavioural measurements discussed in this article. For a more comprehensive definition of each measure, see STAR Methods.

## Results

### Variation in social behaviour

The movement of some species, such as *D. mojavensis, D. immigrans*, *D. virilis*, *D. santomea* and *D. mauritiana*, were noticeably sedentary, compared to more active species such as *D. melanogaster* (Figure 2). Overall, the male flies in all species were more active than the female flies, except for *D. novamexicana* and the outgroup species *Chymomyza procnemis* (Figure 2). In *D. yakuba*, *D. erecta* and *D. hydei*, the males were markedly more active than their female counterparts. Through a two-way ANOVA, we found significant species-by-sex interaction effects for movement (p < 0.0001, Figure 2).

**Figure 2:**
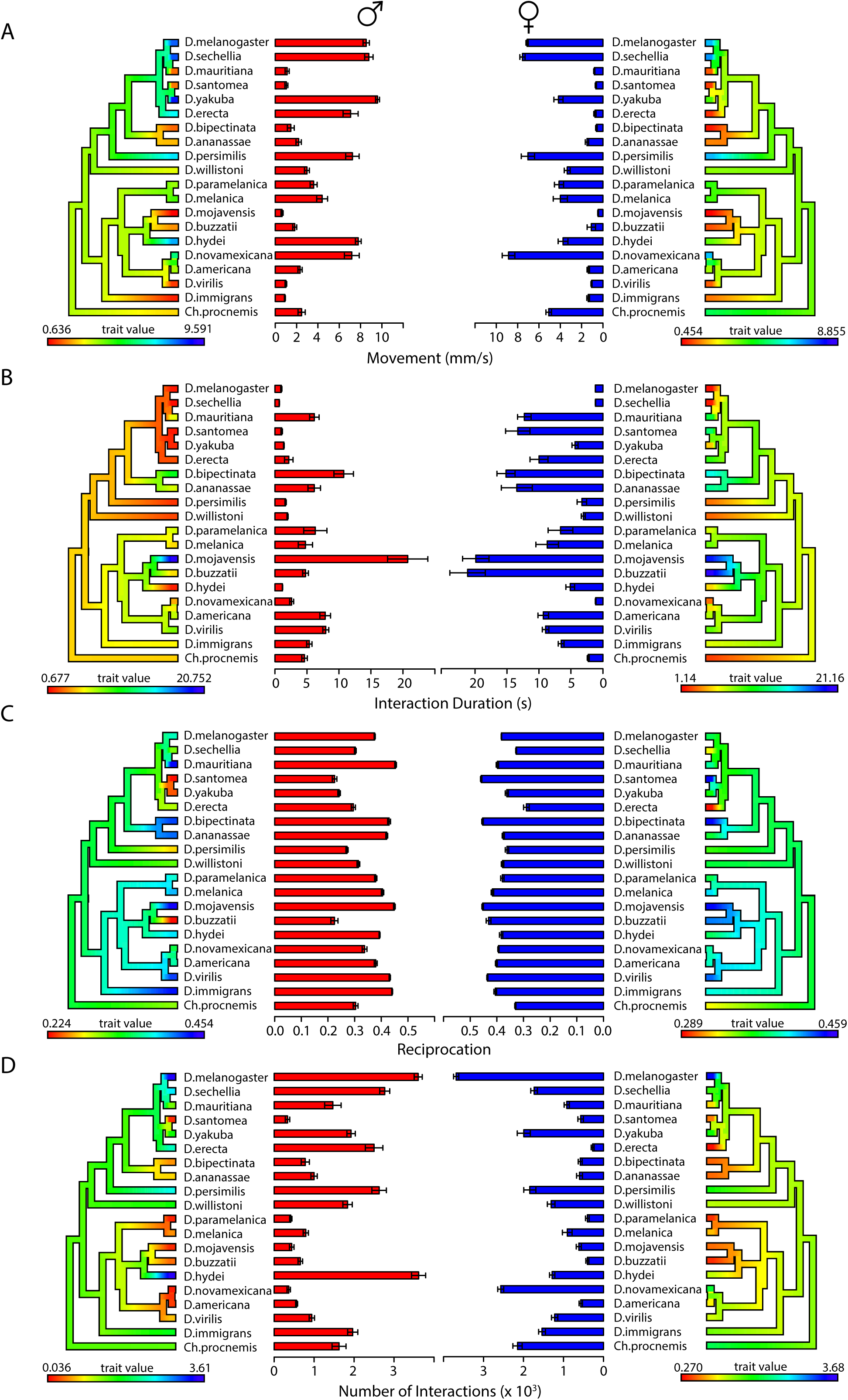
Movement, interaction duration, reciprocation and number of interactions vary across species and between groups of male flies and female flies. The y-axis lists each species, along with a phylogeny mapping their evolutionary relationships. Each phylogeny is coloured using the ‘contMap’ function (phytools package), according to a scale where red indicates the minimum mean measure observed and blue indicates the maximum mean measure observed. Error bars indicate standard error of the mean. All graphs on the left with red bars refer to the data acquired from groups of male flies and all graphs on the right with blue bars refers to the data acquired from groups of female flies. A) Movement: Significant species-specific effects were observed (p < 2 x 10^-16^, two-way ANOVA). Significant sex specific effects and sex by species interaction effects were observed (p = 2 x 10^-16^ for both effects; two-way ANOVA). B) Interaction duration: Significant species-specific effects were observed (p < 2 x 10^-16^, two-way ANOVA). Significant sex specific effects and sex by species interaction effects were observed (p = 2 x 10^-16^ for both effects; two-way ANOVA). C) Reciprocation: Significant species-specific effects were observed (p < 2 x 10^-16^, two-way ANOVA). Significant sex specific effects and sex by species interaction effects were observed (p = 2 x 10^-16^ for both effects; two-way ANOVA). D) Number of interactions: Significant species-specific effects were observed (p < 2.24 x 10^-11^, two-way ANOVA). Significant sex specific effects and sex by species interaction effects were observed (p = 2 x 10^-16^ for both effects; two-way ANOVA).

Next, we observed differences in the social spacing parameters across the drosophilid species in both the male and female data sets (Table S2). The distance and time parameter displayed the least variation for both the male dataset (distance: 1.25-3.25 body lengths; time: 0.2-2.65 seconds) and the female dataset (distance: 1.25-2.75 body lengths; time: 0.35-1.85 seconds). In the male dataset, the species with the estimated distance parameter of 3.25 body lengths was the outgroup species *Ch. procnemis*. Thus, in the male dataset, all *Drosophila* species’ distance parameter ranged from 1.25 to 2.50 body lengths (Table S2). The angle parameter showed the most variation in both the male dataset (20-147.5 degrees) and the female dataset (30-140 degrees). Most female species interact at a wider angle than their male counterparts. This is especially evident in *D. santomea*, *D. yakuba*, *D. willistoni*, *D. persimilis* and *D. buzzatii*, where the females have an estimated angle parameter nearly double the parameter of the males (Table S2). Overall, this data shows that most males and females of the same species interact differently and that most females interact at a wider angle than the males of the same species.

When we accounted for the different social spacing parameters, we found that there were significant species-by-sex interaction effects for interaction duration, reciprocation and number of interactions (p < 0.0001 for all measures, two-way ANOVA, Figure 2). For reciprocation, female flies tended to reciprocate interactions more frequently than male flies. This may be attributable to the wider angle parameters estimated in the female social spacing data (Table S2). For interaction duration, there is a clear inverse relationship with movement, especially in the female dataset where sedentary species spent more time socially interacting on average. Species that were more active also interacted more on average. Although some sedentary species interacted less than active species, all species interacted at least hundreds of times on average (Figure 2), providing relatively large samples of SINs to analyze. Taken together, these data suggest that species vary in the characteristics of their pairwise social interactions.

Despite the variation in movement, social spacing and pairwise interaction patterns, all 20 species form SINs in at least 80% of the video trials in both the male and female datasets (Figure S1). We found significant species-by-sex interaction effects for assortativity (p = 0.0007), clustering coefficient (p < 0.0001), betweeness centrality (p < 0.0001), and global efficiency (p < 0.0001; two-way ANOVA). Qualitatively across all SIN measures, the relative species differences appear very consistent between the male and female datasets (Figure 3): clustering coefficient and global efficiency display the largest range of species’ variation, compared to betweeness centrality and assortativity (Figure 3). Prior to collecting the 20 species-wide data, we collected data for seven species. Five common species were compared between these two independent data sets. The estimated SIN measures (Figures S5-S6) were consistent between these independent data collections. Overall, these data suggest that SIN structure varies across species and sex and that species’ SIN structures are stable across time.

**Figure 3:**
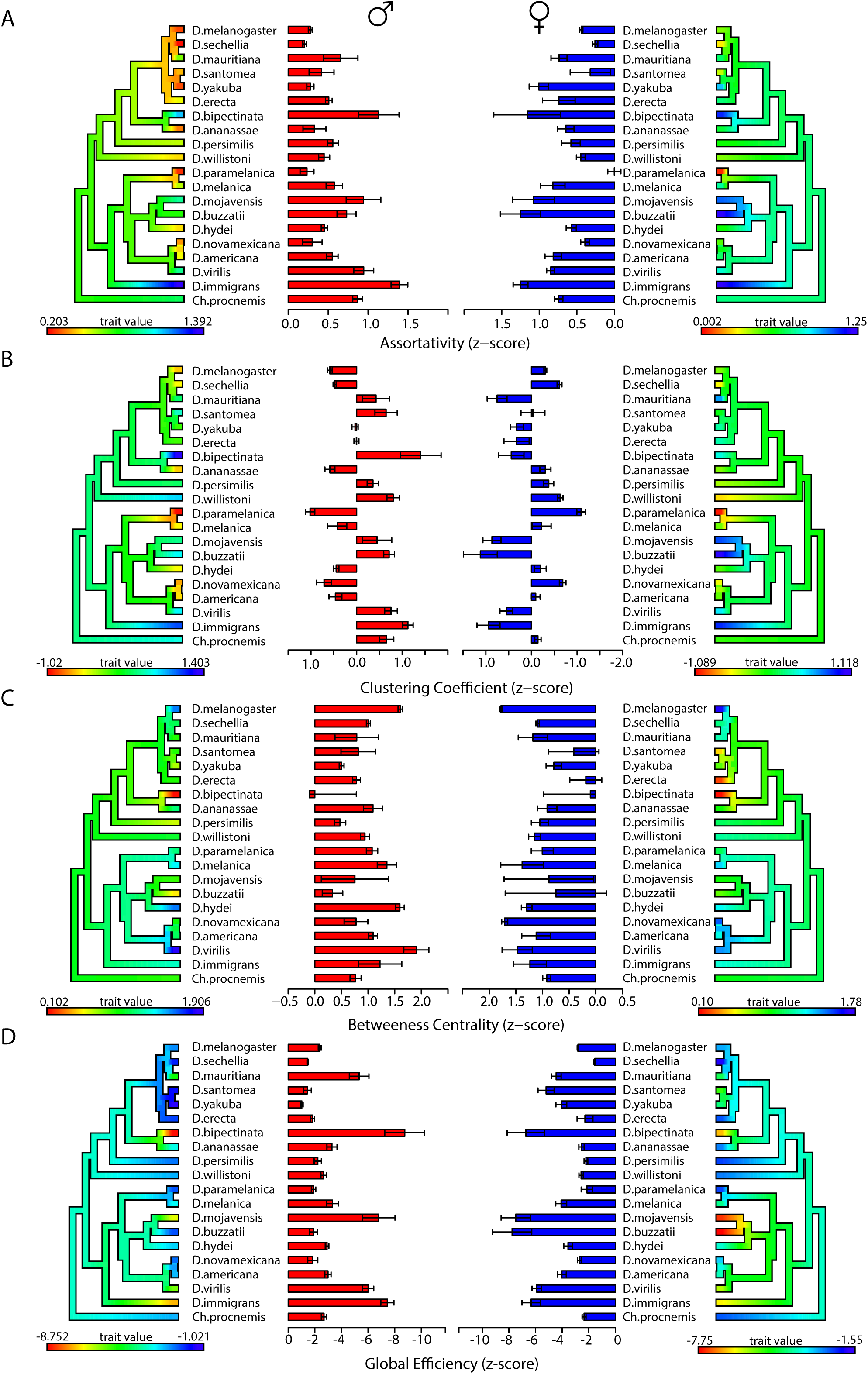
SINs vary across species and between groups of male flies and female flies. For each measure, the x-axis is expressed as a z-score, normalized from the values of virtual SINs (see Methods). The y-axis lists each species, along with a phylogeny mapping their evolutionary relationships. Each phylogeny is coloured using the ‘contMap’ function (phytools package), according to a scale where red indicates the minimum mean measure observed and blue indicates the maximum mean measure observed. Error bars indicate standard error of the mean. All graphs on the left with red bars refer to the data acquired from groups of male flies and all graphs on the right with blue bars refers to the data acquired from groups of female flies. A) Assortativity: Significant species-specific effects were observed (p < 2 x 10^-16^, two-way ANOVA). Significant sex-specific and sex by species interaction effects were observed (p = 0.0002, p = 0.0007 respectively, two-way ANOVA). B) Clustering coefficient: Significant species-specific effects were observed (p < 2 x 10^-16^, two-way ANOVA). No significant sex specific effects were observed (p = 0.248, two-way ANOVA) but a significant sex by species interaction effect was observed (p < 2 x 10^-16^, two-way ANOVA). C) Betweeness centrality: Significant species-specific effects are observed (p < 2 x 10^-16^, two-way ANOVA). No sex-specific effects were observed following multiple test correction (p = 0.0151, two-way ANOVA). A significant sex by species interaction effect was observed (p = 7.65 x 10^-15^, two-way ANOVA). D) Global efficiency: Significant species specific, sex specific, and sex by species interaction effects were observed (p = 7.23 x 10^-9^, p < 2 x 10^-16^, p < 2 x 10^-16^ respectively; two-way ANOVA). See also Figures S1 for the percentage of SINs formed across all video trials and Figures S5-S6 for replication of SIN measures from pilot data.

### Phylogenetic signal

To test for phylogenetic influence on comparative trait data, two metrics were implemented: Blomberg’s K [24] and Pagel’s λ [25]. High phylogenetic signal implies traits are conserved within a given phylogenetic tree. Low phylogenetic signal can be explained by rapidly changing evolutionary events and we favor the explanation that the trait is conserved beyond the root of the given phylogenetic tree. We applied these phylogenetic signal tests across all the behavioural elements (movement, social spacing, pairwise interactions and SIN variables). We found the strongest evidence of high phylogenetic signal for the social spacing distance parameter in both the male and female datasets (Table 2). Both the K and λ tests agree on this statistically significant result, except for the K value in the male dataset (K = 0.20, p = 0.21). This disagreement between the two tests may result from statistical noise because a sample size of 20 species is considered minimal to detect phylogenetic signal [24, 27].

**Table 2:**
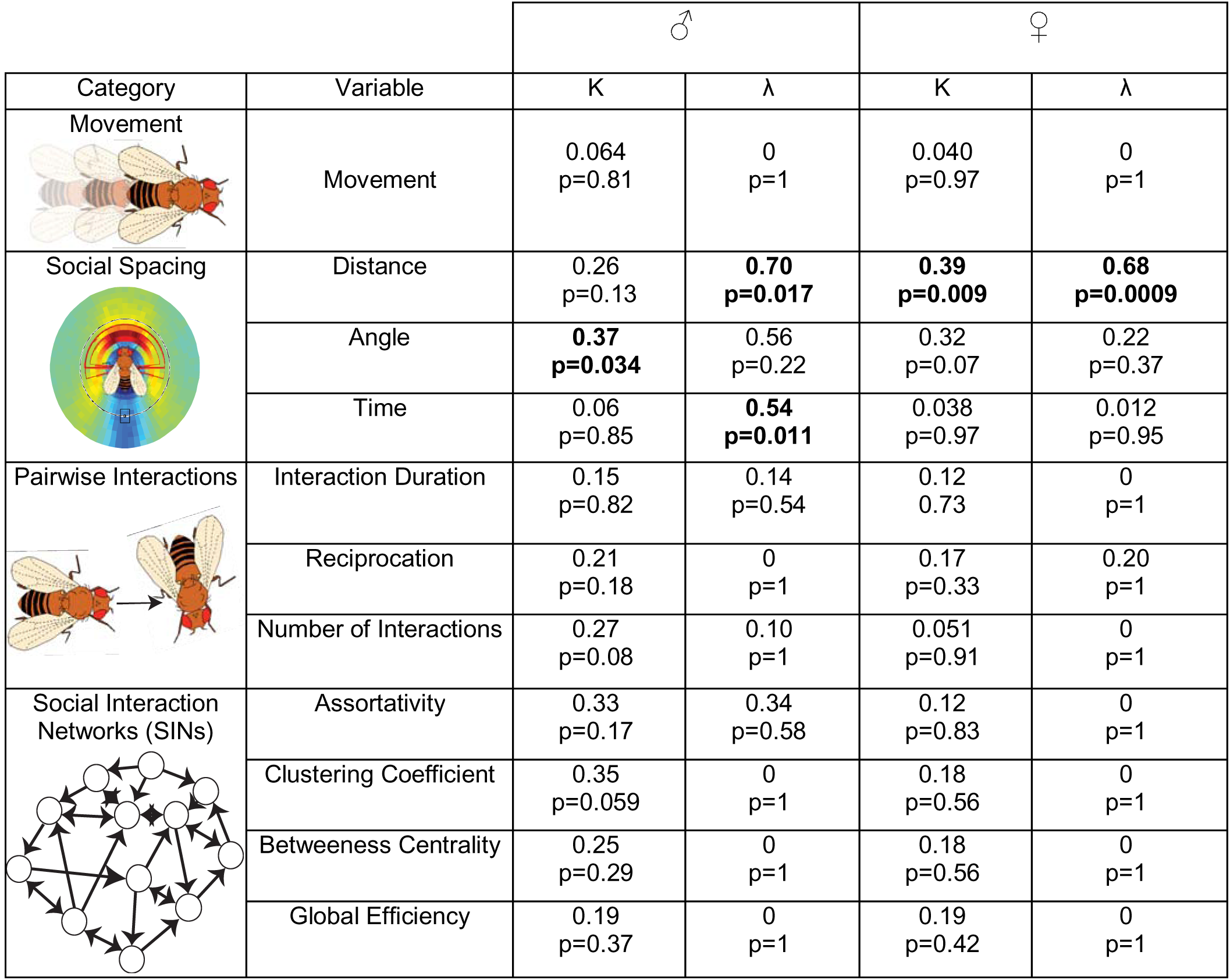
Phylogenetic signal test results of all behavioural variables for K and λ Bolded cells indicate statistical significance for the presence of phylogenetic signal.

| Category | Variable | ♂ |  | ♀ |  |
| --- | --- | --- | --- | --- | --- |
| | | K | $\lambda$ | K | $\lambda$ |
| Movement | Movement | 0.064<br>p=0.81 | 0<br>p=1 | 0.040<br>p=0.97 | 0<br>p=1 |
| Social Spacing | Distance | 0.26<br>p=0.13 | **0.70**<br>**p=0.017** | **0.39**<br>**p=0.009** | **0.68**<br>**p=0.0009** |
|  | Angle | **0.37**<br>**p=0.034** | 0.56<br>p=0.22 | 0.32<br>p=0.07 | 0.22<br>p=0.37 |
|  | Time | 0.06<br>p=0.85 | **0.54**<br>**p=0.011** | 0.038<br>p=0.97 | 0.012<br>p=0.95 |
| Pairwise Interactions | Interaction Duration | 0.15<br>p=0.82 | 0.14<br>p=0.54 | 0.12<br>0.73 | 0<br>p=1 |
|  | Reciprocation | 0.21<br>p=0.18 | 0<br>p=1 | 0.17<br>p=0.33 | 0.20<br>p=1 |
|  | Number of Interactions | 0.27<br>p=0.08 | 0.10<br>p=1 | 0.051<br>p=0.91 | 0<br>p=1 |
| Social Interaction Networks (SINs) | Assortativity | 0.33<br>p=0.17 | 0.34<br>p=0.58 | 0.12<br>p=0.83 | 0<br>p=1 |
|  | Clustering Coefficient | 0.35<br>p=0.059 | 0<br>p=1 | 0.18<br>p=0.56 | 0<br>p=1 |
|  | Betweenness Centrality | 0.25<br>p=0.29 | 0<br>p=1 | 0.18<br>p=0.56 | 0<br>p=1 |
|  | Global Efficiency | 0.19<br>p=0.37 | 0<br>p=1 | 0.19<br>p=0.42 | 0<br>p=1 |

To illustrate the phylogenetic signal for the distance parameter, hereafter referred to as social distance, we mapped each species’ distance parameter (Table S2) onto the drosophilid phylogeny. This produced an evolutionary trait map that showed strong phenotypic divergence of social distance between the *Drosophila* and *Sophophora* subgenera (Figure 4). We hypothesized that this divergence in social distance between the two subgenera may correlate with differences in leg size since: 1) most of the *Drosophila* species in our sample were noticeably larger than the *Sophophora* species; 2) all species commonly extended their legs to touch conspecifics during social interactions. To test this, we measured the legs and bodies of male flies for all 20 species. To control for the differences in body sizes across all species, we calculated a relative leg length measure (total leg length ÷ body size; Table S3 and S4). We generated a trait map of relative leg length which shows that the *Sophophora* subgenus has a longer social distance with longer relative leg lengths and the *Drosophila* subgenus has a shorter social distance with shorter relative leg lengths (Figure 4). Next, we tested whether the relative leg length predicts social distance within the entire *Drosophila* genus by generating a simple linear regression model (Figure 4). These results show a positive correlation (R^2^=0.259) between relative leg length and social distance.

**Figure 4:**
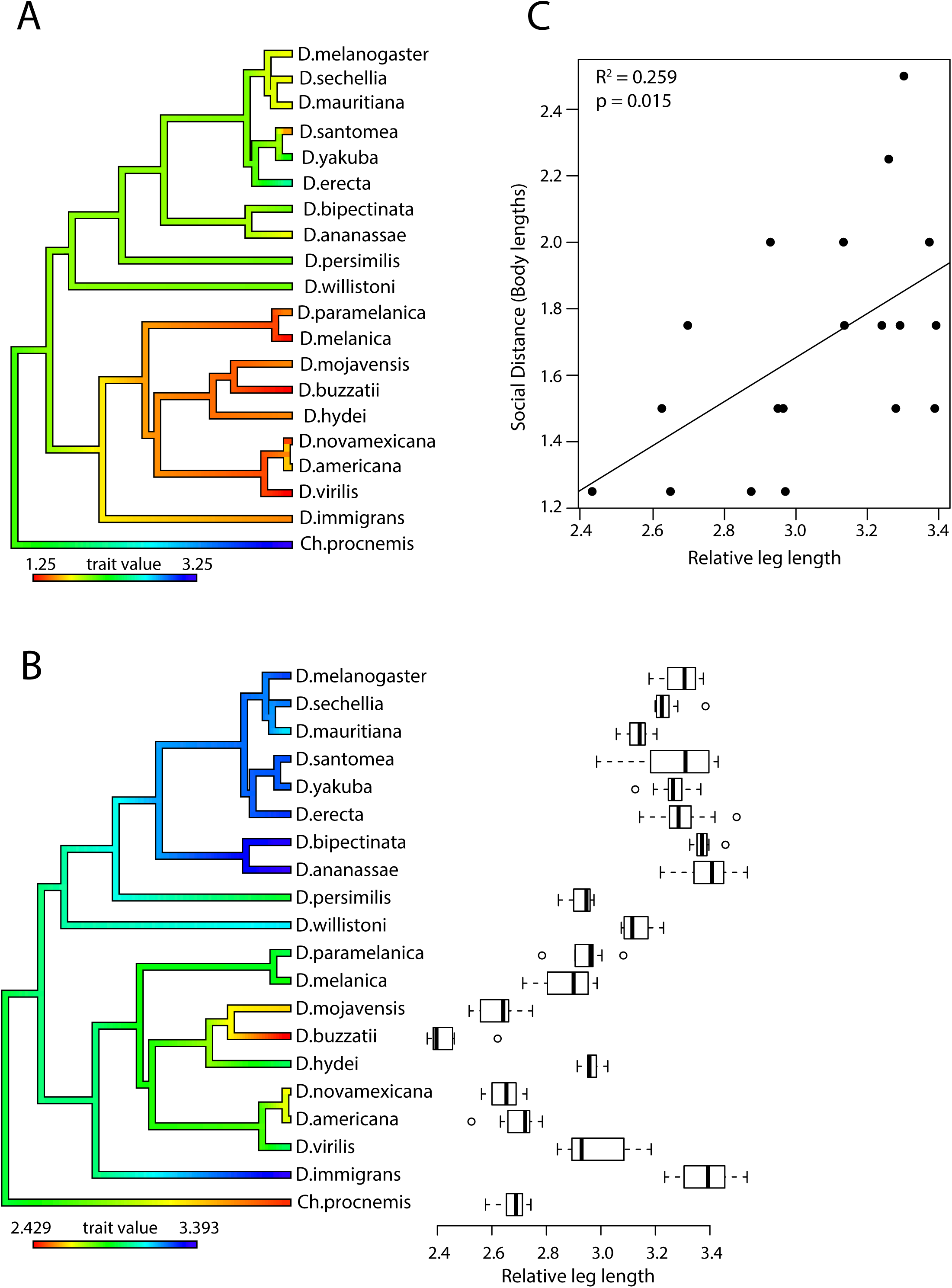
Social distance and relative leg length display phylogenetic signal and are correlated. A) Phylogenetic trait map of drosophilid species differences in social distance. The phylogeny is coloured using the ‘contMap’ function (phytools package) to help visualize species differences and ancestral nodes. Red indicates the minimum value and blue indicates the maximum value observed across the species. The social distance displays divergence between the *Drosophila* and *Sophophora* subgenera. B) Boxplot displaying drosophilid species differences across the relative leg length (total leg length / relative body size). Circles depict outliers, whiskers represent the maximum and minimum values and the bolded line within each box represents the median. The phylogeny on the y-axis is coloured using the ‘contMap’ function (phytools). C) Simple linear regression outlining the relationship between relative leg length (x-axis) and social distance (y-axis). See also Tables S3-S4 for measurements of leg length and body size across all species.

To sum up, these data show that most properties of self-organization in drosophilids have low phylogenetic signal, except social distance which has strong phylogenetic signal. We also show the physical characteristics, leg length and body size, correlates with social distance in *Drosophila*.

### Environmental models of social behaviour

To investigate whether past environmental selective pressures predict current social structure across drosophilids, we extracted 19 climate variables from WorldClim [26]. We simplified the 19 climate variables to 5 principal components and used them as predictors to generate environmental models, for each SIN measure, through stepwise regressions. Temperature range, precipitation of the wettest quarter and temperature of the coldest quarter were strong proponents in each of the climatic principal components (data not shown). We found the resultant environmental models to be surprisingly predictive for assortativity (R^2^ = 0.444), clustering coefficient (R^2^ = 0.452), betweeness centrality (R^2^ = 0.314), and global efficiency (R^2^ = 0.352; Figure 5). We generated additional environmental models via stepwise regression by fitting the 5 principal components to the behavioural element variables. The subsequent environmental models were less predictive than the models for SIN measures: movement (R^2^ = 0.250), the social spacing parameters (distance: R^2^ = 0.255, angle: R^2^ = 0.145, time: R^2^ = 0.160) and the pairwise interaction variables (number of interactions: R^2^ = 0.208, interaction duration: R^2^ = 0.382, reciprocation: R^2^ = 0.122; Figure S2-S4). Together this indicates that climatic selective pressures correlate to group-level behaviours better than they correlate to the individual behavioural elements.

**Figure 5:**
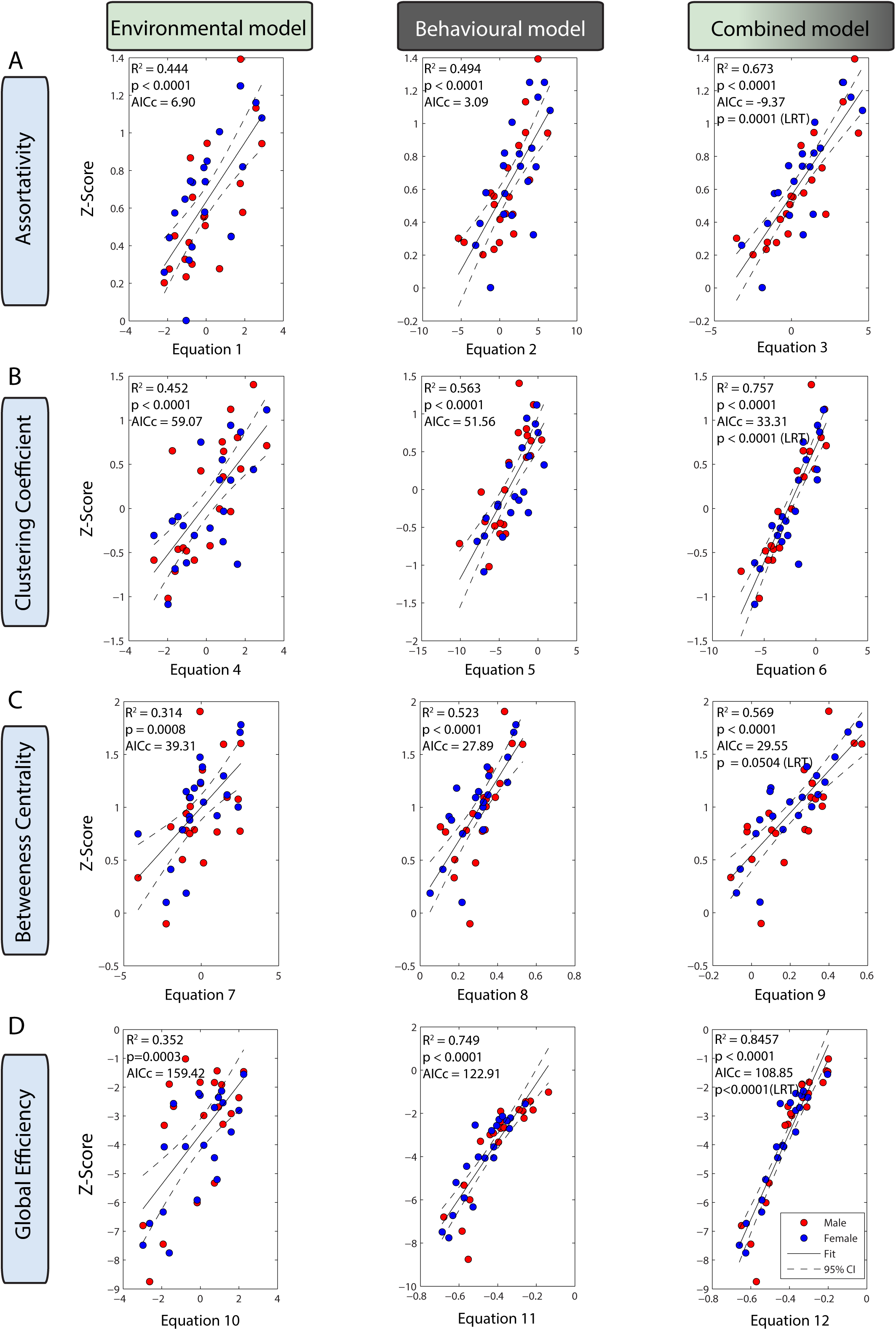
Environmental, behavioural and combined stepwise regression models of assortativity, clustering coefficient, betweeness centrality and global efficiency. For all regressions, each data point represents the mean SIN measure for a single species. The mean SIN measure for groups of male flies and female flies were pooled into each regression and are labeled with red and blue points, respectively. Each solid trend line indicates line of best fit and dashed lines indicate 95% confidence interval of the model. For each model, the multivariate linear equation can be found below. A) The environmental model (Equation 1) significantly predicts assortativity (R^2^ = 0.44447, p < 0.0001). The behavioural model (Equation 2) significantly predicts assortativity (R^2^ = 0.49493, p < 0.0001). The combined model (Equation 3) significantly improves the prediction of assortativity compared to behavioural model alone (Combined model AICc < Behavioural model AICc; p = 0.0001, log likelihood ratio test). B) The environmental model (Equation 4) significantly predicts clustering coefficient (R^2^ = 0.45258. p < 0.0001). The behavioural model (Equation 5) significantly predicts clustering coefficient (R^2^ = 0.56293, p < 0.0001). The combined model (Equation 6) significantly improves the prediction of clustering coefficient compared to behavioural model alone (Combined model AICc < Behavioural model AICc; p < 0.0001, log likelihood ratio test). C) The environmental model (Equation 7) significantly predicts betweeness centrality (R^2^ = 0.31432, p= 0.0008). The behavioural model (Equation 8) significantly predicts betweeness centrality (R^2^ = 0.52325, p < 0.0001). The combined model (Equation 9) does not significantly improve the prediction of betweeness centrality compared to behavioural model alone (Combined model AICc > Behavioural model AICc; p = 0.0504, log likelihood ratio test). D) The environmental model (Equation 10) significantly predicts global efficiency (R^2^ = 0.35261, p = 0.0003). The behavioural model (Equation 11) significantly predicts global efficiency (R^2^ = 0.74972). The combined model (Equation 12) significantly improves prediction of global efficiency (Combined AICc < Behavioural AICc; p < 0.0001, log likelihood ratio test). Equation 1: y = 0.63 – 0.092*[PC2] + 0.064*[PC3] + 0.11*[PC5]. Equation 2: y = 0.53 – 0.081*[movement] + 0.023*[interaction duration] + 0.00018*[number of interactions]. Equation 3: y = 0.56 – 0.068*[PC2] + 0.045*[PC3] + 0.086*[PC5] – 0.071*[movement] + 0.11 *[interaction duration] + 0.00019*[number of interactions]. Equation 4: y = 0.047 – 0.069*[PC1] – 0.18*[PC2] + 0.21*[PC5]. Equation 5: y = 0.73 – 0.19*[movement] – 0.0072*[angle] + 0.04*[interaction duration] + 0.00038*[number of interactions]. Equation 6: y = 0.71 – 0.014*[PC1] – 0.15*[PC2] + 0.14*[PC5] – 0.16*[movement] – 0.0067*[angle] + 0.021*[duration] + 0.00038*[number of interactions]. Equation 7: y = 0.99 + 0.07*[PC1] + 0.057*[PC2] – 0.13*[PC5]. Equation 8: y = 0.096 – 0.28*[social distance] + 0.25*[time] – 0.031*[interaction duration] + 0.0002*[number of interactions] + 2.91*[reciprocation]. Equation 9: y = 0.54 + 0.0065*[PC5] + 0.055*[PC2] – 0.11*[PC5] – 0.39*[social distance] + 0.23*[time] – 0.01*[interaction duration] + 0.0002*[number of interactions] + 1.94*[reciprocation]. Equation 10: y = −3.64 + 0.55[PC2] – 0.34[PC3] – 0.58[PC5]. Equation 11: y = 2.03 + 0.31*[movement] – 0.14*[interaction duration] – 0.0007*[number of interactions] – 13*[reciprocation]. Equation 12: y = 2.47 + 0.33*[PC3] – 0.38*[PC5] + 0.26*[movement] – 0.078*[interaction duration] – 0.0007*[number of interactions] – 15*[reciprocation]. See also Figures S2-S4 for environmental and behavioural models of other behavioural elements.

### Behavioural models of social behaviour

To investigate how behavioural elements predict SINs, we fit all the behavioural element variables (movement, social spacing, pairwise interactions) to each SIN measure through stepwise regressions. We found the resultant behavioural models to be strongly predictive for assortativity (R^2^ = 0.495), clustering coefficient (R^2^ = 0.587), betweeness centrality (R^2^ = 0.523) and global efficiency (R^2^ = 0.760; Figure 5). Interestingly, movement contributed to all behavioural models for SIN measures, except betweeness centrality (Figure 5). We decided to investigate our behavioural hierarchy by correlating higher-order behaviours to their respective lower-order behaviours (Figure 1; see Methods for explanation of lower- and higher-order behaviours). We found these behavioural models to be strongly predictive (number of interactions: R^2^ = 0.538, interaction duration: R^2^ = 0.553, and reciprocation: R^2^ = 0.879; Figure S3-S4). Together, this suggests that more complex higher-order behaviours in *Drosophila* can be predicted by some lower-order behaviours but not all lower-order behaviours (such as movement) can predict complex social structures.

### Combined models of social behaviour

We tested how much variation in each SIN measure can be explained by both the environmental and behavioural models simultaneously in a “combined model”. We hypothesized that comparing the combined model to the behavioural model through a likelihood ratio test would determine whether the combination of the two models outperforms the behavioural model alone. We found that the combined models significantly improve the fit over the behavioural models for assortativity (R^2^ = 0.673, p = 0.0001; likelihood ratio test), clustering coefficient (R^2^ = 0.757, p < 0.0001) and global efficiency (R^2^ = 0.846, p < 0.0001; Figure 5). The combined model for betweeness centrality remains nearly identical to the behavioural model (Combined: R^2^ = 0.569, Behavioural: R^2^ = 0.523), suggesting that the behavioural data alone is sufficient at predicting betweeness centrality. We repeated this analysis by generating combined models for the pairwise interaction variables (level 3; interaction duration, number of interactions and reciprocation). We found that the combined models do not significantly outperform the behavioural models for any of these variables (Figure S4), indicating that movement (level one) and social space variables (level two) are sufficient at predicting pairwise interactions (level three). Altogether, this suggests that the best prediction of SIN measures requires the consideration of both environmental and behavioural factors. However, betweeness centrality and pairwise interactions can be sufficiently predicted by behaviour alone.

## Discussion

In this study, we set out to determine which factors shape social organization by studying social networks in *Drosophila*. To do this, we used a species-wide comparative method to model climate and behaviour to the evolution of SINs. Here, we show that all 20 drosophilid species used in this study form SINs (Figure S1) and that the properties of these SINs vary across species (Figure 3). This suggests that all drosophilids self-organize but not all drosophilids self-organize similarly. We show the same results from a pilot experiment we performed, suggesting we are capturing a reproducible, stable phenotype (Figure S5-S6). We found low phylogenetic signal for the measured SIN properties and speculate that group organization in drosophilids is the result of conserved mechanisms stemming deep in evolutionary time.

We found that social distance, which represents the typical distance at which flies communicate, displays evidence of high phylogenetic signal according to Pagel’s λin males and females and Blomberg’s K in females (Table 2). According to our data, most species tend to interact within 1-3 body lengths, and this range of interaction distance has been reported and applied by other researchers [5, 13, 28–30]. The distance at which flies socially interact is conserved within different clades of our tree, suggesting that closely related drosophilids evolved similar mechanisms for social communication (Figure 4). We hypothesized that the social distance at which flies interact may correlate with the physical limitation of a fly’s anatomical ability to extend its legs towards another fly, which contain gustatory receptors [31]. In flies, it is thought that touching conspecifics leads to the exchange of pheromones [32, 33], which can serve as cues about a flies’ social environment [34, 35]. It is also possible that flies communicate with each other using both touch and taste. We show that relative leg length does positively correlate with social distance (Figure 4). This suggests that how flies space themselves in a social setting may be determined by the length of their legs relative to their body size and we postulate that their leg length determines how close they need to be to conspecifics to touch. Evidence of flies touching has been documented in *D. paramelanica* [36], *D. melanogaster* [13] and witnessed in this study across all species. We reason that social interactions within a species may influence the evolution of their anatomy, such as leg length. By maintaining the ability to interact and communicate with conspecifics, individuals are better equipped to survey their environment [34, 35], locate potential mates and find food [2]. The evolution of leg length and social distance in *Drosophila* may be one example of this.

We found that the past climatic pressures, behavioural elements, and a combination of these variables can predict SIN structure in drosophilids. Although the behavioural elements we investigate in this study are also behaviours that comprise SINs, they are not necessarily predictive of SIN structure. For example, different sensory mutants in *D. melanogaster* show significantly different movement and rates of interaction. However, the sensory mutants form SINs with the same clustering coefficient, assortativity, and betweeness centrality [13]. In this study, we show that different combinations of behavioural elements can be used to predict SIN structure across different *Drosophilia* species.

We found that past climatic pressures correlate to the behavioural elements that comprise drosophilid SINs and to the different properties of their SINs (Figure S2-S4, Figure 5, respectively). These climatic variables could be indicative of other past pressures of their environment, such as food availability [37, 38], species richness [39] and intraspecific density [40]. Thus, we argue that the ways in which drosophilids behave and the structures of their groups have evolved, at least in part, by pressures exerted by their environments. It has been shown that how individuals organize in groups can influence their fitness and health [3–5]. Thus, understanding the influences on social organization is imperative for recognizing that disrupting these influences can impact the survival of a species. It has been speculated that 40% of the insect populations on earth will become extinct in the next few decades and that climate change will be one factor contributing to their destruction [41]. However, not all declining insect populations can be attributable to climate change alone and, in some cases, factors contributing to the observed differences in insect populations remain unexplained [41, 42]. Seeing as social organization has been shown to contribute to health and reproductive success, it is rational to speculate that the breakdown of social organization, brought on by climate change, is contributing to the extinction of these insects.

In this study, we found that movement varies across drosophilid species. Interestingly, the most sedentary species in this study tend to organize themselves into SINs with high clustering coefficient (Figure 3). This could lead to a social structure where flies interact more equally [43]. Some studies suggest that living in more egalitarian groups correlates with longer lifespans [44] and, interestingly, some of the most sedentary species in our study, such as *D. mojavensis* and *D. virilis*, have been reported to have some of the longest life spans across the *Drosophila* genus [45]. We argue that the differences in group organization seen across drosophilid species is influenced by their different geographic origins and that these differences may correlate with life span. In the future, we may be able to evaluate properties of social networks and predict health outcomes of populations, such as longevity.

Although movement correlated with three SIN measures, it did not correlate with betweeness centrality (Figure 5), a SIN measure previously reported to be heritable in *Drosophila* [13] and humans [15]. Moreover, when we combined the environmental and behavioural models for betweeness centrality, the combined model did not significantly improve the correlation compared to the behavioural model alone (Figure 5). It appears that the climate data adds no value to maximizing the prediction of betweeness centrality, and that the behavioural elements are sufficient. We speculate that betweeness centrality, as a phenotype, may respond primarily to selective pressures of the social environment. In one study, researchers found that the willingness of individuals to share or withhold information from competitors is influenced by the composition of the social environment, which they coined “audience effects” [46]. In *Drosophila*, audience effects may explain why flies signal fertile oviposition sites [6, 7] and the presence of predators to naïve flies [10, 11]. In the wild, flies may readily organize and transmit valuable information about their environment if the benefits of cooperation exceed the cost of competition. This may facilitate dynamic behavioural strategies which vary across individuals, populations and species. Overall, our results suggest that betweeness centrality may measure a group-level phenotype that is likely genetically heritable and has been selected by pressures of the social environment. Future network studies on *Drosophila*, or in other animals, may be able to determine gene(s) responsible for regulating the betweeness centrality phenotype, which could lead to the discovery of conserved social mechanisms that may be found in other organisms.

We highlight how phylogeny, environment and simple behaviours influence the social organization of drosophilids. All 20 of the drosophilids we studied socially organize and form SINs, allowing for the possibility of dissecting the evolution of sociality in the future. We speculate that social communication across drosophilids is conserved, given that drosophilids are known to encounter and interact with other species [47]. Recent work uncovered that drosophilids can socially signal the presence of predators to an unrelated heterospecific species [11], suggesting that flies communicate in “dialects” with each other. When considering complex social organization behaviours, the ecological pressures of a species’ environment play a large role in shaping these phenotypes. Perhaps, one day, genes that influence social organization will be uncovered in *Drosophila* and we will consider social organization as a behavioural phenotype that emerges from a deeply conserved genetic toolkit [48].

## Supporting information

Supplementary Material Jezovit et al

## Acknowledgements

Funding for this study was graciously provided to JDL by NSERC, CIHR, the Canada Research Chair, and Canadian Institute for Advanced Research. The authors wish to thank Leslie Vosshall, Michael Dickinson, John Ratcliffe, Rob Ness, Marc Johnson, and Luke Mahler for helpful discussions along the way.

## Author Contributions

J.A.J. acquired the data, analyzed the data and wrote the first draft of this manuscript. R.R. assisted with some experiments and participated in editing and writing the final draft of this manuscript. J.S. wrote some of the scripts for data analysis, provided statistical insights and participated in data analysis. J.A.J., J.S., J.D.L. designed the experiments and edited the manuscript.

## STAR⍰Methods

### KEY RESOURCES TABLE

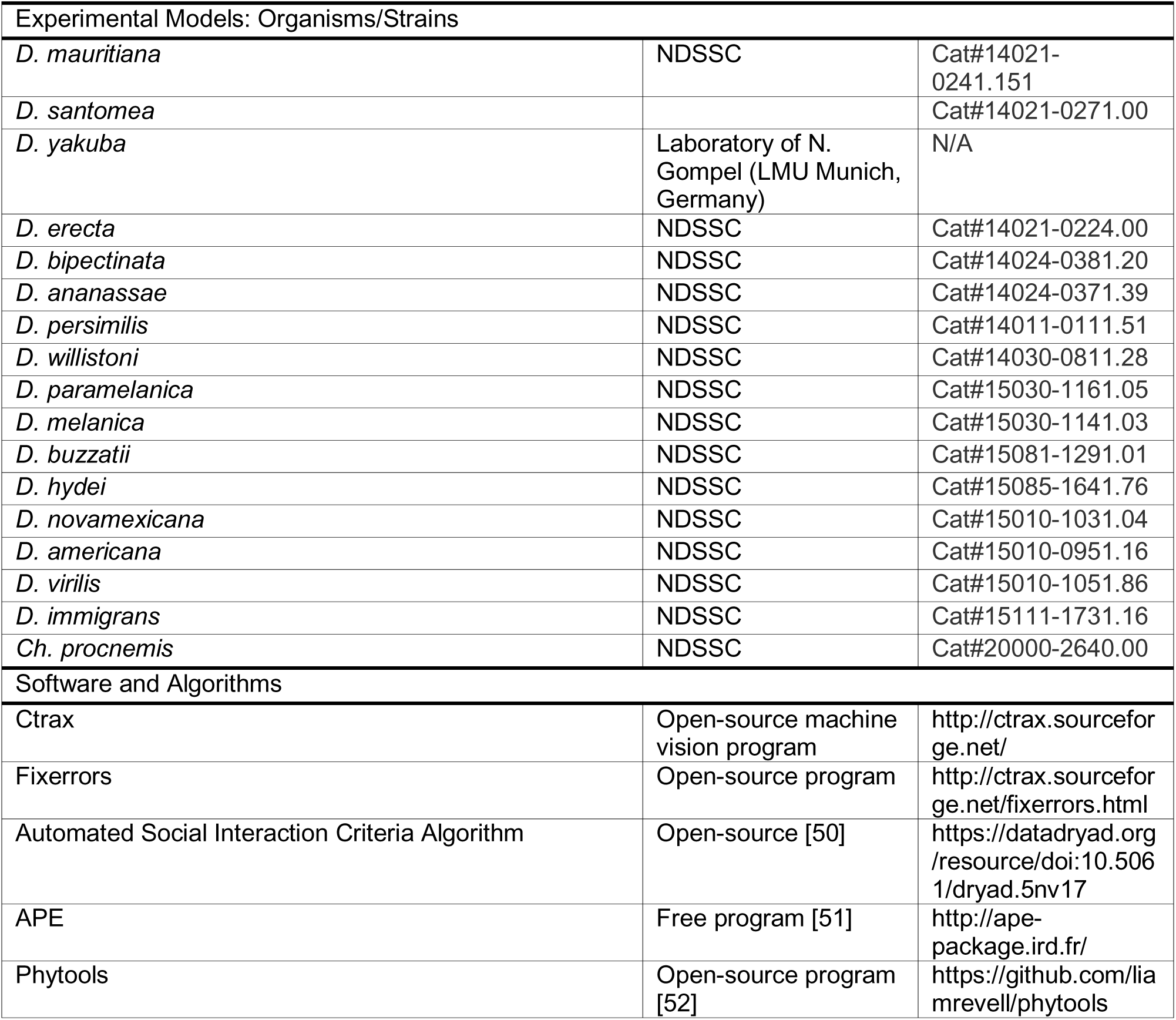

### LEAD CONTACT AND MATERIALS AVAILABILITY

Further information and requests for resources and reagents should be directed to and will be fulfilled by the lead contact Joel Levine.

This study did not generate new unique reagents.

### EXPERIMENTAL MODEL AND SUBJECT DETAILS

#### Fly stocks

The majority of the species used for the SIN assay were purchased from the *Drosophila* Species Stock Centre (http://blogs.cornell.edu/drosophila/orders/) as shown in Table S1. The Canton-S strain was used to represent *D. melanogaster*. The *D. yakuba* stock was a gift from N. Gompel. The source of the *D. mojavensis* and *D. sechellia* stocks are unknown, but these stocks were confirmed to be the correct species through documented phenotypic markers [17].

### METHOD DETAILS

#### Video acquisition

SIN data were acquired as outlined previously [13]. To summarize, all fly stocks were maintained in bottles with 40 mL of media containing cornmeal, wheat germ, soy flour, molasses, sucrose, glucose, yeast, agar, propionic acid and Tegosept. These bottles were stored in a Percival incubator (model I-36LL) set to 25°C with a 12 h L:D cycle. For all species, virgin male and female flies were collected through light CO_2_ anaesthesia. Male and female flies were housed separately in groups of 12-16 individuals within vials containing 8 mL of media. All flies were reared for 3 days in an incubator controlled under the same temperature and photoperiod described above. Afterwards, 12 flies from each vial were gently aspirated into circular arenas (60 mm diameter, 2 mm depth) and filmed for 30 minutes using Firefly MV cameras (Point Grey). All videos were filmed at the same time of day (9.5-10.5 h after initiation of light phase during photoperiod) to control for the clock-controlled locomotor activity of flies. All filming was completed inside a Biochamber with controlled temperature (25°C) and humidity (65% RH). After filming, all flies were anaesthetized with CO_2_ and discarded. The position, orientation and identity of all flies in each video were tracked through machine vision software (Ctrax; versions 0.4.2 & 0.5.19b). Errors in tracking were manually corrected through the fixerrors (version 0.2.3) MATLAB package to correct for: swapping of fly identities, errors in the orientation of the head to abdomen region of the fly tracks and drastic fluctuations in the large major axis of the fly tracks. Data collection was performed separately for groups of male flies and for groups of female flies and we will use the terms “male dataset” and “female dataset”, respectively. In total, we collected, tracked and fixed the errors in 1000 video trials, totaling approximately forty million frames of video, across the male and female datasets of the 20 drosophilid species.

#### Pilot data

An additional male and female dataset containing 399 videos across 7 species was collected and analyzed. This collection of videos was acquired between July 2014 and August 2015. The following species are represented in this pilot data: *D. melanogaster* (Canton-S), *D. simulans*, *D. sechellia*, *D. yakuba*, *D. pseudoobscura*, *D. mojavensis*, and *D. paramelanica*. With this pilot data, we were able to test the robustness of SIN measures between 5 species common to both independent data collections. The stocks for these species were the same as those used in the 20 species-wide dataset. The relative species differences, for each dataset, were statistically tested through a Kruskal-Wallis one-way ANOVA in MATLAB. The replication of these datasets were visualized through boxplots that display the distribution of the SIN scores for each species (Figure S5-S6).

#### Automated estimation of social spacing interaction criteria

All interaction criteria used to generate SINs in the 20 species-wide data (referred to as extended data below) are shown in Table S2. The distance, angle and time interaction criteria were estimated for each species using an algorithm published by Schneider and Levine [50]. Stated briefly, the algorithm analyzes the spatial positions of every fly in all tracked videos. The algorithm eliminates background noise from flies by analyzing spatial positions of “virtual trials” which consist of fly tracks randomly sampled from separate videos. With that background subtraction, the algorithm identifies distance, angle and time criteria that are over-represented in videos of flies socially interacting compared to the non-social virtual trials. Initially, the algorithm provided interaction criteria that contradicted personal observations of each species’ spatial patterning. The following adjustments were made to improve the performance of the algorithm:

1. Previously the algorithm computed an “inter-fly distance” defined as the interval where the distribution for distance measures was positive when subtracted from the virtual trials. This interval was the starting point to locate interaction hotspots on the social space heat map (see S1-S25 for a visualization of these heat maps). Here, we did not compute the inter-fly distance and established the initial interaction hotspots by targeting the 95^th^ percentile instead of the third quartile. As reported previously [50], angle and distance bins were increased until the mean of the enclosed region on the heat maps began to decrease.
2. Previously the algorithm computed the time criteria by recording the time elapsed when flies fulfilled the calculated distance and angle criteria. Background noise would be removed by subtracting the distribution of time values from the virtual trial time distribution. The first positive time bin, representing the minimal time value over-represented in videos of socially interacting flies, was utilized as the time criteria. Here, the time criteria were acquired by increasing the number of bins on the normalized time frequency distribution until the mean of the enclosed bins decreases.

For each species, this algorithm was performed 500 times where 15 videos were randomly sampled with replacement in order to generate a 95% confidence interval and median estimate of each species’ interaction criteria. Both the pilot data (see below) and extended data were analyzed with this algorithm. For a few species, the estimated interaction criteria produced questionable results that contradicted our own observations. The following changes were made to amend the computerized estimates:

1. *D. mauritiana* males: the angle criterion was relaxed to 121.5 degrees since that angle encompasses more hotspots in their social space heat map (data not shown).
2. *D. santomea* females: the distance was restricted to the lower limit of the 95% confidence interval, the angle was relaxed to the upper limit of the 95% confidence interval, and time was restricted to the lower limit of the 95% confidence interval (data not shown).
3. *D. immigrans* males: The automated time criterion was estimated to be 0 s, clearly a failure of the algorithm to provide a meaningful criterion. As a result, 103 social interactions of male *D. immigrans* were randomly flagged throughout the videos and the duration of these interactions (in units of seconds) was recorded in a histogram. The minimum value of 0.53 s was used to generate SINs. This value falls in a bin that contains ∼ 10% of the interactions observed, making it a reliable approximation of a minimum social interaction time criterion (data not shown).
4. *D. mojavensis* males and females: the angle criterion computed in the extended data was narrower than the angle criterion estimated from the pilot data. Because the pilot data angle criterion encompasses a larger distribution of hot spots that are consistent with the extended data heat maps, the pilot data angle criterion was used for SIN analysis in the extended data (Table S2; data not shown).

#### Estimation of social interaction networks and other behavioural measures

We considered that different drosophilid species may interact differently. We considered how the inter-individual distance, angle and time, collectively referred to as the social spacing parameters (Figure S2), may vary across species. To account for this, we utilized an algorithm that estimates these parameters based on tracked videos (see above). Each species’ distance, angle and time parameters for both the male and female datasets are listed in Table S2. These species-specific and sex-specific social spacing parameters allowed us to control for variation in social interactions when generating SINs. Additionally, the social spacing parameters can also be used as a measure to classify the ways that different species interact.

All SINs were generated as previously described [13]. To summarize, all SINs are iterative networks that comprise 33 unique interactions, which represents a network density of 25% (25% of 132 unique possible interactions in a group of 12 individuals). The number of iterative networks in a single 30-minute video may vary and is proportional to the number of unique interactions in a video. Most species formed at least one SIN iteration in over 80% of their respective video trials (Figure S1). We assess four SIN measures that we view as social organization phenotypes: i) *assortativity*, defined as the probability of an individual interacting with another individual with a similar degree (degree is defined as the number of incoming and outgoing connections to a single node); ii) *clustering coefficient*, defined as a measure of how interconnected neighbours are to one another; iii) *betweeness centrality*, defined as the number of shortest paths that traverse an individual, indicating the relative importance of an individual for maintaining the cohesion of the network; and iv) *global efficiency*, defined as a measure of redundant pathways, indicating the efficiency of information flow throughout the network [13, 14]. Explanations of these SIN measures that correspond to the iterative approach are described in Table 1. Here we speak about these network measures as they apply to a group on average. The four SIN measures are expressed as z-scores, which normalizes the networks to control for degree distribution. To do this, we generated 10000 random networks and calculated the z-score as follows:

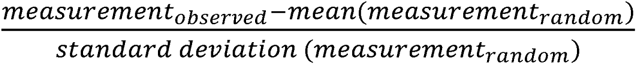

For each SIN measure, we present the mean z-score, averaged from the distribution of videos for each species in the male and female dataset (Figure 3; Schneider et al., 2012 methods [13]).

In addition to the SIN measures, other behavioural elements were estimated from the tracked videos: i) *movement*, defined as the mean locomotor activity (millimetres/second) of the flies in a single video; ii) *interaction duration*, defined as the mean duration of the social interactions (based on the distance, angle and time parameters) in a single video; iii) *reciprocation*, defined as the proportion of social interactions reciprocated in a single video; and iv) *number of interactions*, defined as the total number of social interactions in a single video. See Table 1 for a summary of these behavioural elements. Like the SIN measures, the mean of each behavioural element is calculated from the distribution of videos for each species in the male and female dataset (Figure 2).

#### Phylogenetic signal

To determine whether the drosophilid species’ social behaviour exhibit phylogenetic signal (i.e. differences in a given trait reflect the species’ degree of common ancestry), we applied two tests: Blomberg’s K [24] and Pagel’s λ [25]. Both phylogenetic signal tests evaluate the extent to which the observed average trait data across species adhere to a model of Brownian motion across a phylogenetic tree, which approximates the expectation under evolution due to random genetic drift [53, 54]. Both the K and λ values typically range on a scale from 0-1, where 0 indicates low phylogenetic signal (trait evolution does not adhere to Brownian motion) and 1 indicates high phylogenetic signal (trait evolution adheres to Brownian motion; see Blomberg et al. 2003 [24]). The phylogenetic tree applied to these analyses was a published drosophilid phylogeny that we pruned to include our species sample [21]. Since *D. novamexicana* was not included in this tree, its placement and branch length was standardized based on another published phylogenetic tree of the *virilis* group species [55]. The tree was made ultrametric in R through the ‘chronos’ function (ape package) for all subsequent phylogenetic comparative analyses.

#### Geographic coordinate acquisition for the 20 species

To test the influence of climatic environment on species variation in SINs, climate data was acquired from each species geographic distribution. Many of the species purchased from the *Drosophila* Species Stock Centre had records of the geographic coordinates, and/or the city and country the stock was collected from. For the stocks that did not have precise coordinates listed on the stock center, the rough latitude and longitude of the recorded locations were noted from a search on Google Maps. The coordinates estimating each species’ stock geographic origin are listed in Table S1. However, we were unsure of the precise geographic origin for the following stocks: *D. sechellia*, *D. yakuba*, *D. erecta*, *D. mojavensis*. As a result, we utilized Taxodros, a database that contains records of coordinates pin-pointing thousands of collection sites for drosophilid species (http://www.taxodros.uzh.ch/). We acquired all the Taxodros coordinates listed for each of our 20 species through custom MATLAB scripts. For the species stocks that had known coordinates from the stock centre or Google Maps, we filtered out all Taxodros coordinates beyond +/- 2 latitudinal and longitudinal units from the known coordinates (Table S1). Fortunately, the species without known coordinates (*D. mojavensis*, *D. yakuba*, *D. erecta*, *D. sechellia*), have narrow geographic distributions confined to one continent. Therefore, the mean latitude and longitude were calculated from all Taxodros coordinates to approximate the center of their geographic distribution, and these coordinates are listed in Table S1.

The Taxodros database also aided in acquiring more accurate estimates of the climate variables used in the environmental models (Figure 5). Rather than estimating a single measure for all 19 climate variables using the coordinates in Table S1, we acquired distributions for the 19 climate variables from the filtered Taxodros coordinates. For all species, the mean value was calculated from the distribution of each climate variable, and these mean measures were incorporated into the principal component analysis. The first 5 principal components accounted for 92% of the variance across all the climate data (Figure S7).

#### Estimation of environmental variables

We estimated climate data from latitudinal and longitudinal coordinates specific to the capture site of each drosophilid species stock (Figure 1, Table S1). For each set of coordinates, we obtained 19 climate variables from the WorldClim database (http://www.worldclim.org; [26]). All climate variables represent the predicted climate from the mid-Holocene period, approximately 3000 to 8000 years ago [56]. We emphasize that this climate data best captures past selective pressures that each drosophilid species adapted to. The following variables, representing temperature and precipitation patterns, were obtained: *annual mean temperature* (BIO1)*, mean diurnal range* (BIO2; calculated as mean of monthly(max temp – min temp)), *isothermability* (BIO3; calculated as BIO2/BIO7 * 100), *temperature seasonality* (BIO4), *maximum temperature of warmest month* (BIO5), *minimum temperature of coldest month* (BIO6), *temperature annual range* (BIO7; calculated as BIO5-BIO6), *mean temperature of wettest quarter* (BIO8), *mean temperature of driest quarter* (BIO9), *mean temperature of warmest quarter* (BIO10), *mean temperature of coldest quarter* (BIO11), *annual precipitation* (BIO12), *precipitation of wettest month* (BIO13), *precipitation of driest month* (BIO14), *precipitation seasonality* (BIO15), *precipitation of wettest quarter* (BIO16), *precipitation of driest quarter* (BIO 17), *precipitation of warmest quarter* (BIO 18), *precipitation of coldest quarter* (BIO 19).

#### Leg measurements

The front, middle and rear legs were measured from a minimum of 10 individuals for all 20 drosophilid species (Table S3). First, we anaesthetized the flies with carbon dioxide and carefully severed each leg above the femur using forceps. Each leg was placed on a flat surface and covered with a coverslip. An image was acquired for each leg using a Zeiss SteREO Discovery.V12 microscope and Zeiss ZEN (2011) software. The leg was measured in µm using Zeiss ZEN (2011) built-in measurement tools. The femur measurement began where the trochanter met the femur and ended at the tibia. The tibia was measured from the beginning of the tibia to the end of the tibia. The tarsi were measured from the beginning of the tarsi and ended in the middle of the claw. For each leg, the length of the femur, tibia and tarsal fragments were summed. The sum of the three legs were then averaged to provide us with the total leg length.

#### Body size measurements

The body sizes for each species were measured using the tracked videos (see *Video acquisition* above). For each 30-minute tracked video, the mean large major axis was calculated in µm for each fly across all frames. Then, the mean of all individual flies within each video was calculated. Therefore, each video generated a single mean body size measurement. Each species’ reported body size (Table S4) was calculated by averaging the mean body size measurements from all video trials. Since the body sizes varied for each species, we calculated the relative leg length by dividing the total leg length by body size.

### QUANTIFICATION AND STATISTICAL ANALYSIS

#### SINs and behavioural elements

Social interaction network (SIN) data was acquired from 19 *Drosophila* species and one outgroup species, *Chymomyza procnemis*. We gathered data for 20 species since a power analysis reported this as a reliable minimum sample size for phylogenetic comparative methods [24, 27]. Approximately 20-25 videos were acquired for each species in both the male and female datasets (see Figure S1 for precise sample sizes of each species).

A two-way ANOVA was used in MATLAB to test whether SINs and the behavioural elements differ by sex and across species. Due to the repeated tests on 8 measures, a Bonferroni correction was employed and all p-values below 0.00625 were considered significant. Statistical details can be found in the results, figures and figure legends.

Outliers were removed from all analyses. For the SIN measures and behavioural measures, a data was considered an outlier if it was greater than the 75^th^ quartile+(1.5*IQR) or lower than the 25^th^ quartile-(1.5*IQR).

#### Phylogenetic signal

The K and λ tests were implemented in R using the ‘phylosig’ function (phytools package). We tested phylogenetic signal on the four SIN measures and the four behavioural elements where each species’ mean and standard error of the mean were incorporated to represent the species’ average and intraspecific variation for the trait, respectively [27]. We also tested phylogenetic signal for the distance, angle and time social spacing parameters. Since some of these variables were manually altered, we did not incorporate any measure of intraspecific variation into the phylogenetic signal tests for these measures. All phylogenetic signal tests were performed separately on the male and female data sets. The estimated K values were statistically evaluated using R through permutation tests where the observed K value was compared to a null distribution of 1000 K values, each computed by randomly shuffling the tips of the phylogenetic tree. All estimated λ values were statistically evaluated using R through likelihood ratio tests against the null hypothesis that λ = 0. All measures of phylogenetic signal and the associated p-values are present in Table 2. Statistical details can be found in the results, figures and figure legends. Each phylogenetic signal test had a sample size of 20, equal to the number of species. All p-values < 0.05 were considered statistically significant.

To visualize the phylogenetic signal tests for each variable, trait data was mapped onto the drosophilid phylogeny using the ‘contMap’ function (phytools package) in R. This function maps ancestral states for the internal nodes of the phylogeny using maximum likelihood, given the trait data for each species at the tips of the tree [57]. These phylogenetic trait maps are present in Figures 2-4.

#### Regression analyses

To explore how environment and behavioural elements influence drosophilid species’ variation in SINs, we produced three different types of models through multivariate stepwise regressions: i) environmental models, ii) behavioural models and iii) combined models (Figure 5). The environmental models were derived from the mid-Holocene climatic data described above. Through a principal component analysis implemented in MATLAB, we reduced the 19 climate variables to 5 principal components that account for 92% of the observed variance in the climate data (Figure S7). For each environmental model, the 5 principal components served as the initial predictors, with one SIN measure as a response variable. To generate each environmental model, the “stepwiselm” function, was implemented in MATLAB with default parameters. The optimal predictors that remained after forward and backward stepwise selection are listed below the x-axis of each regression (Figure 5).

The behavioural models were produced through stepwise regressions, similar to the environmental models. The following variables served as predictors and were fit to each SIN measure: movement, social spacing parameters (distance, angle and time) and the pairwise interaction variables (interaction duration, reciprocation, number of interactions; Figure 1). The purpose of these models was to explore how behavioural elements predict the complex SIN measures. Like the environmental models, the predictors that remained after forward and backward selection are listed below the x-axis of each regression (Figure 5).

Finally, the combined models were generated by combining and fitting the predictors of the associated environmental and behavioural models to each SIN measure (Figure 5). This was done to test if the environmental data, together with the behavioural data, improves the correlation to each SIN measure. To test this, we compared the fit of the combined model to the behavioural model for each SIN measure through a likelihood ratio test. We justify that the comparison can be done against only the behavioural models because their R^2^ value was consistently better than the environmental models for each SIN measure. Significant results indicate the combination of behavioural and environmental variables are better at modeling the SIN measures than either the behavioural variables or environmental variables alone. In total, 10 variables had behavioural and combined models generated and compared (see Figure 5, Figures S2-S4). Therefore, we considered α ≤ 0.005 as significant through a Bonferroni correction. In addition, we also validated the quality of the combined models, relative to the behavioural models, by comparing their Akaike Information Criterion (AICc) values.

We characterized the behavioural elements in a hierarchy, where movement is level one, social spacing is level two, pairwise interactions are level three, and SINs are level four (Figure 1). Separate environmental models, behavioural models and combined models were generated using level two and three variables as response variables. To generate level three behavioural models, level one and two variables (movement, social spacing) served as predictors. To generate level two behavioural models, the level one variable (movement) served as a predictor. Both male and female data were pooled in these regression analyses since the SIN and behavioural measures contain significant sex by species interaction effects (Figure 2-3). All data points in the regression are the mean values for each behavioural measure.

To explore whether total leg length and relative leg length predicts the level two distance parameter (social distance), we generated simple linear regressions in R. For each regression we excluded the outgroup, *Ch. procnemis*, since we were interested in whether social distance correlated with the total and relative leg lengths of species belonging to the *Drosophila* genus.

### DATA AND CODE AVAILABILITY

The behavioural videos supporting the current study have not been deposited in a public repository because of the immense size of the ∼1000 videos but are available from the corresponding author on request.

## Supplemental Information

**Table S1-Related to Method Details: Summary of species stocks used for all experiments.** All stocks purchased from the *Drosophila* Species Stock Centre are indicated. Where applicable, the geographic origin of the capture site of each stock is indicated. The approximate geographic coordinates of each stock are also indicated, and these were utilized for acquisition of climate data.

**Table S2-Related to Method Details: Interaction criteria of all 20 species for both male and female flies.** Interaction criteria in bold are values that were used to over-ride the automated estimations (see methods).

**Table S3-Related to Figure 4: Leg lengths of all 20 species for male flies.** Mean front, middle, and rear legs were measured and a total mean leg length was calculated.

**Table S4-Related to Figure 4: Mean body size of all 20 species for male flies.** Body sizes were calculated from tracked videos and averaged for each species.

**Figure S1-Related to Method Details and Figure 3: Ability to form SINs for the male dataset (blue) and female dataset (red).** The x-axis is expressed as a percentage of the number of videos with at least one SIN iteration divided by the total number of videos acquired for each species. There are no differences across species in their ability to form networks (p = 1; χ goodness of fit test). The total number of videos that formed SINs are as follows: *D. melanogaster*: n= 46 (male), n=48 (female); *D.sechellia*: n=22 (male), n=22 (female); *D. mauritiana*: n=26 (male), n=27 (female); *D. santomea*: n=21 (male), n=23 (female); *D. yakuba*: n=24 (male), n=26 (female); *D.erecta*: n=23 (male), n=21 (female); *D. bipectinata*: n=22 (male), n=24 (female); *D. ananassae*: n=23 (male), n=25 (female); *D. persimilis*: n=23 (male), n=25 (female); *D. willistoni*: n=23 (male), n=24 (female); *D.paramelanica*: n=22 (male), n=18 (female); *D. melanica*: n=22 (male), n=21 (female); *D. mojavensis*: n=25 (male), n=25 (female); *D. buzzatii*: n=23 (male), n=21 (female); *D. hydei*: n=25 (male), n=24 (female); *D. novamexicana*: n=24 (male), n=22 (female); *D. virilis*: n=25 (male), n=27 (female); *D. immigrans*: n=23 (male), n=22 (female); *Ch. procnemis*: n=20 (male), n=21 (female).

**Figure S2-Related to Figure 1: Environmental model for the level 1 variable Movement.** For the regression, each data point represents the mean SIN measure for a single species. The mean SIN measure for groups of male flies and female flies were pooled into each regression and are labeled with red and blue points, respectively. The solid trend line indicates line of best fit and dashed lines indicate 95% confidence interval of the model. The equation for this model is listed below the x-axis.

**Figure S3-Related to Figure 1: Environmental, behavioural and combined models for the level 2 social spacing variables: Distance, Angle and Time.** For all regressions, each data point represents the mean SIN measure for a single species. The mean SIN measure for groups of male flies and female flies were pooled into each regression and are labeled with red and blue points, respectively. Each solid trend line indicates line of best fit and dashed lines indicate 95% confidence interval of the model. **A)** No significant behavioural model formed for distance, therefore no combined model to report. The environmental model (Equation 1 predicts social distance (R^2^ = 0.255, p = 0.00164). **B)** No significant behavioural model formed for angle, therefore no combined model to report. The environmental model (Equation 2) predicts angle (R^2^ = 0.145, p = 0.0087). **C)** The environmental model (Equation 3) predicts time (R^2^ = 0.16, p = 0.0062). The behavioural model (Equation 4) predicts time (R^2^ = 0.0837, p = 0.039). The combined model significantly improves prediction of time compared to the behavioural model alone (Combined AICc < Behavioural AICc; p = 0.0003, likelihood ratio test). **Equation 1:** y = 1.80 – 0.071*[PC1] – 0.13*[PC5]. **Equation 2:** y = 106.3 + 4.71*[PC1]. **Equation 3:** y = 0.99 + 0.077*[PC1]. **Equation 4:** y = 1.21 – 0.061*[movement]. **Equation 5:** y = 1.28 + 0.091*[PC1] – 0.078*[movement].

**Figure S4-Related to Figure 1: Environmental, behavioural and combined models for the level 3 variables interaction duration (Interaction** Δ**t), number of interactions (Interaction #), and reciprocation.** For all regressions, each data point represents the mean SIN measure for a single species. The mean SIN measure for groups of male flies and female flies were pooled into each regression and are labeled with red and blue points, respectively. Each solid trend line indicates line of best fit and dashed lines indicate 95% confidence interval of the model. **A)** The environmental model (Equation 1) predicts interaction duration (R^2^ = 0.382, p = 0.0003). The behavioural model (Equation 2) predicts interaction duration (R^2^ = 0.553, p < 0.0001). The combined model (Equation 3) does not significantly improve the prediction of interaction duration compared to the behavioural model alone (Combined AICc =∼ Behavioural AICc; p = 0.0182, likelihood ratio test). **B)** The environmental model (Equation 4) predicts number of interactions (R^2^ = 0.20772, p = 0.005). The behavioural model (Equation 5) predicts number of interactions (R^2^ = 0.538, p < 0.0001). The combined model does not significantly improve the prediction of number of interactions compared to the behavioural model alone (Combined AICc =∼ Behavioural AICc; p = 0.1594, likelihood ratio test). **C)** The environmental model (Equation 7) predicts reciprocation (R^2^ = 0.122, p = 0.034). The behavioural model (Equation 8) predicts reciprocation (R^2^ = 0.879, p < 0.0001). The combined model (Equation 9) does not significantly improve the prediction of reciprocation compared to the behavioural model alone (Combined AICc =∼ Behavioural AICc; p = 0.153, likelihood ratio test). **Equation 1:** y = 8.17 – 0.76*[PC2] + 1.02*[PC3] + 2.31*[PC5]. **Equation 2:** y = 6.54 – 1.25*[movement] + 0.045*[angle]. **Equation 3:** y = 6.99 – 0.23*[PC2] + 0.47*[PC3] + 1.47*[PC5] – 0.93*[movement] + 0.038*[angle]. **Equation 4:** y = 1398 + 220*[PC4] – 336*[PC5]. **Equation 5:** y = 1001 + 213*[movement] – 406*[time]. **Equation 6:** y = 1100 + 91.5*[PC4] – 151*[PC5] + 183*[movement] – 387*[time]. **Equation 7:** y = 0.39 + 0.011*[PC3]. **Equation 8:** y = 0.21 – 0.0065*[movement] + 0.014*[social distance] + 0.0017*[angle] – 0.025*[time]. **Equation 9:** y = 0.22 + 0.0028*[PC3] – 0.0061*[movement] + 0.01*[social distance] + 0.0017*[angle] – 0.025*[time].

**Figure S5-Related to Figure 3: Relative species differences between *D. melanogaster* (CS), *D. sechellia* (SEC), *D. yakuba* (YAK), *D. mojavensis* (MOV), *D. paramelanica* (PARA) are replicated in male flies between pilot data (Experiment 1; left panels) and extended data (Experiment 2; right panels).** All figures are boxplots which outline the distribution of the z-scores for all four SIN measures. Letters above each box indicate statistically distinct groups from a Kruskal-Wallis one-way ANOVA followed Tukey-Kramer post-hoc tests.

**Figure S6-Related to Figure 3: Relative species differences between *D. melanogaster* (CS), *D. sechellia* (SEC), *D. yakuba* (YAK), *D. mojavensis* (MOV), *D. paramelanica* (PARA) are replicated in female flies between pilot data (Experiment 1; left panels) and extended data (Experiment 2; right panels).** All figures are boxplots which outline the distribution of the z-scores for all four SIN measures. Letters above each box indicate statistically distinct groups from a Kruskal-Wallis one-way ANOVA followed Tukey-Kramer post-hoc tests.

**Figure S7-Related to Figure 5: The first 5 principal components of the climatic measures, extracted from each species geographic origin, account for 92% of the variance across the 19 variables.** Each bar represents the percentage of variance explained by the first 10 dimensions. Because the rate of decrease reduces after dimension 5, the first 5 dimensions were used to represent the environmental variables for all regression analyses.

## References

1. Scott, J.P. (1956). The analysis of social organization in animals. Ecology 37, 213–221.

2. Krause, J., and Ruxton, G.D. (2002). Living in groups, (Oxford University Press).

3. Drewe, J.A. (2009). Who infects whom? Social networks and tuberculosis transmission in wild meerkats. Proceedings of the Royal Society B: Biological Sciences 277, 633–642.

4. Oh, K.P., and Badyaev, A.V. (2010). Structure of social networks in a passerine bird: consequences for sexual selection and the evolution of mating strategies. The American Naturalist 176, E80–E89.

5. Pasquaretta, C., Battesti, M., Klenschi, E., Bousquet, C.A., Sueur, C., and Mery, F. (2016). How social network structure affects decision-making in Drosophila melanogaster. Proc Biol Sci 283, 20152954.

6. Battesti, M., Moreno, C., Joly, D., and Mery, F. (2012). Spread of social information and dynamics of social transmission within Drosophila groups. Curr Biol 22, 309–313.

7. Sarin, S., and Dukas, R. (2009). Social learning about egg-laying substrates in fruitflies. Proc Biol Sci 276, 4323–4328.

8. Danchin, E., Nöbel, S., Pocheville, A., Dagaeff, A.C., Demay, L., Alphand, M., Ranty-Roby, S., van Renssen, L., Monier, M., Gazagne, E., et al. (2018). Cultural flies: Conformist social learning in fruitflies predicts long-lasting mate-choice traditions. Science 362, 1025–1030.

9. Lihoreau, M., Poissonnier, L.A., Isabel, G., and Dussutour, A. (2016). Drosophila females trade off good nutrition with high-quality oviposition sites when choosing foods. J Exp Biol 219, 2514–2524.

10. Kacsoh, B.Z., Bozler, J., Ramaswami, M., and Bosco, G. (2015). Social communication of predator-induced changes in Drosophila behavior and germ line physiology. Elife 4.

11. Kacsoh, B.Z., Bozler, J., and Bosco, G. (2018). Correction: Drosophila species learn dialects through communal living. PLoS Genet 14, e1007825.

12. Ramdya, P., Lichocki, P., Cruchet, S., Frisch, L., Tse, W., Floreano, D., and Benton, R. (2015). Mechanosensory interactions drive collective behaviour in Drosophila. Nature 519, 233–236.

13. Schneider, J., Dickinson, M.H., and Levine, J.D. (2012). Social structures depend on innate determinants and chemosensory processing in Drosophila. Proc Natl Acad Sci U S A 109 Suppl 2, 17174–17179.

14. Newman, M. (2010). Networks: an introduction, (Oxford University Press).

15. Fowler, J.H., Dawes, C.T., and Christakis, N.A. (2009). Model of genetic variation in human social networks. Proc Natl Acad Sci U S A 106, 1720–1724.

16. Jezovit, J.A., Levine, J.D., and Schneider, J. (2017). Phylogeny, environment and sexual communication across the Drosophila genus. J Exp Biol 220, 42–52.

17. Markow, T.A., and O’Grady, P.M. (2006). Drosophila: A guide to species identification and use, (San Diego, California, USA: Elsevier, Inc).

18. Foster, E.A., Franks, D.W., Morrell, L.J., Balcomb, K.C., Parsons, K.M., van Ginneken, A., and Croft, D.P. (2012). Social network correlates of food availability in an endangered population of killer whales, Orcinus orca. Animal Behaviour 83, 731–736.

19. Kelley, J.L., Morrell, L.J., Inskip, C., Krause, J., and Croft, D.P. (2011). Predation risk shapes social networks in fission-fusion populations. PloS one 6, e24280.

20. Grimaldi, D.A. (1987). Amber fossil Drosophilidae (Diptera), with particular reference to the Hispaniolan taxa. American museum novitates (USA).

21. van der Linde, K., Houle, D., Spicer, G.S., and Steppan, S.J. (2010). A supermatrix-based molecular phylogeny of the family Drosophilidae. Genetics research 92, 25–38.

22. Kellermann, V., Loeschcke, V., Hoffmann, A.A., Kristensen, T.N., Fløjgaard, C., David, J.R., Svenning, J.C., and Overgaard, J. (2012). Phylogenetic constraints in key functional traits behind species’ climate niches: patterns of desiccation and cold resistance across 95 Drosophila species. Evolution 66, 3377–3389.

23. Kellermann, V., Overgaard, J., Hoffmann, A.A., Fløjgaard, C., Svenning, J.C., and Loeschcke, V. (2012). Upper thermal limits of Drosophila are linked to species distributions and strongly constrained phylogenetically. Proc Natl Acad Sci U S A 109, 16228–16233.

24. Blomberg, S.P., Garland, T., and Ives, A.R. (2003). Testing for phylogenetic signal in comparative data: behavioral traits are more labile. Evolution 57, 717–745.

25. Pagel, M. (1999). Inferring the historical patterns of biological evolution. Nature 401, 877–884.

26. Hijmans, R.J., Cameron, S.E., Parra, J.L., Jones, P.G., and Jarvis, A. (2005). Very high resolution interpolated climate surfaces for global land areas. International Journal of Climatology: A Journal of the Royal Meteorological Society 25, 1965–1978.

27. Ives, A.R., Midford, P.E., and Garland Jr, T. (2007). Within-species variation and measurement error in phylogenetic comparative methods. Systematic Biology 56, 252–270.

28. Durisko, Z., Kemp, R., Mubasher, R., and Dukas, R. (2014). Dynamics of social behavior in fruit fly larvae. PLoS One 9, e95495.

29. Lihoreau, M., Clarke, I.M., Buhl, J., Sumpter, D.J., and Simpson, S.J. (2016). Collective selection of food patches in Drosophila. J Exp Biol 219, 668–675.

30. Simon, A.F., Chou, M.T., Salazar, E.D., Nicholson, T., Saini, N., Metchev, S., and Krantz, D.E. (2012). A simple assay to study social behavior in Drosophila: measurement of social space within a group. Genes Brain Behav 11, 243–252.

31. Vosshall, L.B., and Stocker, R.F. (2007). Molecular architecture of smell and taste in Drosophila. Annu Rev Neurosci 30, 505–533.

32. Bontonou, G., and Wicker-Thomas, C. (2014). Sexual Communication in the Drosophila Genus. Insects 5, 439–458.

33. Ferveur, J.F. (2005). Cuticular hydrocarbons: their evolution and roles in Drosophila pheromonal communication. Behav Genet 35, 279–295.

34. Krupp, J.J., Kent, C., Billeter, J.C., Azanchi, R., So, A.K., Schonfeld, J.A., Smith, B.P., Lucas, C., and Levine, J.D. (2008). Social experience modifies pheromone expression and mating behavior in male Drosophila melanogaster. Curr Biol 18, 1373–1383.

35. Billeter, J.C., Atallah, J., Krupp, J.J., Millar, J.G., and Levine, J.D. (2009). Specialized cells tag sexual and species identity in Drosophila melanogaster. Nature 461, 987–991.

36. Sexton, O.J., and Stalker, H.D. (1961). Spacing patterns of female Drosophila paramelanica. Animal Behaviour 9, 77–81.

37. Bergallo, H.G., and Magnusson, W.E. (1999). Effects of climate and food availability on four rodent species in southeastern Brazil. Journal of Mammalogy 80, 472–486.

38. Craine, J.M. (2013). Long-term climate sensitivity of grazer performance: a cross-site study. PLoS One 8, e67065.

39. Currie, D.J., Mittelbach, G.G., Cornell, H.V., Field, R., Guégan, J.F., Hawkins, B.A., Kaufman, D.M., Kerr, J.T., Oberdorff, T., and O’Brien, E. (2004). Predictions and tests of climate-based hypotheses of broad-scale variation in taxonomic richness. Ecology letters 7, 1121–1134.

40. McInerny, M.C., and Cross, T.K. (1999). Effects of lake productivity, climate warming, and intraspecific density on growth and growth patterns of black crappie in southern Minnesota lakes. Journal of Freshwater Ecology 14, 255–264.

41. Sánchez-Bayo, F., and Wyckhuys, K.A. (2019). Worldwide decline of the entomofauna: A review of its drivers. Biological conservation 232, 8–27.

42. Hallmann, C.A., Sorg, M., Jongejans, E., Siepel, H., Hofland, N., Schwan, H., Stenmans, W., Müller, A., Sumser, H., and Hörren, T. (2017). More than 75 percent decline over 27 years in total flying insect biomass in protected areas. PloS one 12, e0185809.

43. Krause, J., James, R., Franks, D.W., and Croft, D.P. (2015). Animal social networks, (Oxford University Press, USA).

44. Barocas, A., Ilany, A., Koren, L., Kam, M., and Geffen, E. (2011). Variance in centrality within rock hyrax social networks predicts adult longevity. PLoS One 6, e22375.

45. Ma, S., Avanesov, A.S., Porter, E., Lee, B.C., Mariotti, M., Zemskaya, N., Guigo, R., Moskalev, A.A., and Gladyshev, V.N. (2018). Comparative transcriptomics across 14 Drosophila species reveals signatures of longevity. Aging cell 17, e12740.

46. Matos, R.J., and Schlupp, I. (2005). Performing in front of an audience: signalers and the social environment. Animal communication networks, 63–83.

47. Markow, T.A. (2015). The secret lives of Drosophila flies. Elife 4.

48. Dickinson, M.H. (2014). Death valley, Drosophila, and the Devonian toolkit. Annual review of entomology 59, 51–72.

49. Peel, M.C., Finlayson, B.L., and McMahon, T.A. (2007). Updated world map of the Köppen-Geiger climate classification. Hydrology and earth system sciences discussions 4, 439–473.

50. Schneider, J., and Levine, J.D. (2014). Automated identification of social interaction criteria in Drosophila melanogaster. Biol Lett 10, 20140749.

51. Paradis, E., and Schliep, K. (2018). ape 5.0: an environment for modern phylogenetics and evolutionary analyses in R. Bioinformatics 35, 526–528.

52. Revell, L.J. (2012). phytools: an R package for phylogenetic comparative biology (and other things). Methods in Ecology and Evolution 3, 217–223.

53. Felsenstein, J. (1985). Phylogenies and the comparative method. The American Naturalist 125, 1–15.

54. Revell, L.J., Harmon, L.J., and Collar, D.C. (2008). Phylogenetic signal, evolutionary process, and rate. Syst Biol 57, 591–601.

55. Spicer, G.S., and Bell, C. (2002). Molecular phylogeny of the Drosophila virilis species group (Diptera: Drosophilidae) inferred from mitochondrial 12S and 16S ribosomal RNA genes. Annals of the Entomological Society of America 95, 156–161.

56. Sandweiss, D.H., Maasch, K.A., and Anderson, D.G. (1999). Transitions in the mid-Holocene. Science 283, 499–500.

57. Revell, L.J. (2013). Two new graphical methods for mapping trait evolution on phylogenies. Methods in Ecology and Evolution 4, 754–759.

