## Supplementary Material Jezovit et al for "The Evolution of Social Organization: Climate Influences Variation in Drosophilid Social Networks"

**Table S1-Related to Method Details: Summary of species stocks used for all experiments**. All stocks purchased from the Drosophila Species Stock Centre are indicated. Where applicable, the geographic origin of the capture site of each stock is indicated. The approximate geographic coordinates of each stock are also indicated, and these were utilized for acquisition of climate data (see supplementary methods).

| **Species** | **Geographic origin** | **GIS coordinates [latitude, longitude]** |
|---|---|---|
| *D. melanogaster* (Canton-S) | Canton, Ohio | [40.46, -81.19] |
| *D. sechellia* | Seychelles islands | [-4.15, 55.45]* |
| *D. mauritiana* | Mauritius | [-20.3484, 57.55215] |
| *D. santomea* | San Tome and Principe Island | [0.23, 6.6] |
| *D. yakuba* | Central-West Africa | [-1.74361, 18.07164]* |
| *D. erecta* | Central-West Africa | [1.863, 3.302]* |
| *D. bipectinata* | Cambodia | [12.56, 104.99] |
| *D. ananassae* | Guam | [13.4443, 144.79373] |
| *D. persimilis* | Santa Cruz Island, California | [37.73, -122.43] |
| *D. willistoni* | Jalisco, Mexico | [20.32,-105.31] |
| *D. paramelanica* | Reedsburg, Wisconsin | [43.5625, -89.8255] |
| *D. melanica* | Austin, Texas | [30.27, -97.74] |
| *D. mojavensis* | Sonora, Mexico | [29.167, -111.947]* |
| *D. buzzatii* | Cochabamba, Bolivia | [-17.41, -66.16] |
| *D. hydei* | Victoria, Australia | [-37.79, 145.43] |
| *D. novamexicana* | Moab, Utah | [38.57, -109.54] |
| *D. americana* | Iowa River, Iowa | [41.96, -93.89] |
| *D. virilis* | Puebla, Mexico | [19.04, -98.20] |
| *D. immigrans* | San Diego, California | [32.84, -117.20] |
| *Ch. procnemis* | Fukuoka, Japan | [33.58, 130.4] |

* Indicates the precise geographic origin of the stock was uncertain

**Table S2-Related to Method Details: Interaction criteria of all 20 species for both male and female flies.** Interaction criteria in bold are values that were used to over-ride the automated estimations (see methods).

| | Male | | | Female | | |
|---|---|---|---|---|---|---|
| **Species** | **Distance** | **Angle** | **Time** | **Distance** | **Angle** | **Time** |
| *D. ananassae* | 1.75 | 140 | 0.95 | 2.25 | 100 | 1.15 |
| *D. bipectinata* | 2 | 127.5 | 1.025 | 1.75 | 130 | 0.7 |
| *D. melanogaster* | 1.75 | 105 | 0.45 | 1.75 | 110 | 0.55 |
| *D. erecta* | 2.5 | 75 | 0.5 | 2.25 | 30 | 0.85 |
| *D. mauritiana* | 1.75 | **121.5** | 0.25 | 2.5 | 95 | 1.25 |
| *D. persimilis* | 2 | 60 | 0.6 | 2 | 110 | 0.85 |
| *D. santomea* | 1.5 | 20 | 0.2 | **2.75** | **135** | **0.4** |
| *D. sechellia* | 1.75 | 70 | 0.35 | 2 | 85 | 0.6 |
| *D. willistoni* | 2 | 70 | 0.6 | 2.25 | 120 | 1.05 |
| *D. yakuba* | 2.25 | 55 | 0.5 | 2.5 | 120 | 1.4 |
| *D. americana* | 1.75 | 130 | 2.65 | 1.5 | 125 | 1.85 |
| *D. buzzatii* | 1.25 | 50 | 0.95 | 1.25 | 115 | 1.3 |
| *D. hydei* | 1.5 | 125 | 0.6 | 1.75 | 135 | 1.3 |
| *D. immigrans* | 1.5 | 125 | **0.53** | 1.5 | 120 | 1.85 |
| *D. melanica* | 1.25 | 135 | 0.9 | 1.25 | 130 | 1 |
| *D. mojavensis* | 1.5 | **147.5** | 1.3 | 1.5 | **140** | 0.35 |
| *D. novamexicana* | 1.25 | 115 | 1.3 | 1.25 | 120 | 0.6 |
| *D. paramelanica* | 1.5 | 130 | 1.75 | 1.5 | 135 | 1.6 |
| *Ch. procnemis* | 3.25 | 35 | 1 | 2.25 | 60 | 0.8 |
| *D. virilis* | 1.25 | 135 | 1.75 | 1.25 | 140 | 1.8 |

**Table S3-Related to Figure 4: Leg lengths of all 20 species for male flies.** Mean front, middle, and rear legs were measured and a total mean leg length was calculated.

| Species | Sample Size | Mean Front Leg (µm) | Mean Middle Leg (µm) | Mean Rear Leg (µm) | Mean Total Leg Length (µm) |
|---|---|---|---|---|---|
| *Ch. procnemis* | 20 | 1808.24 | 2011.56 | 1808.24 | 1876.02 |
| *D. melanogaster* | 10 | 1743.64 | 2184.59 | 2225.42 | 2051.21 |
| *D. sechellia* | 10 | 1509.37 | 1929.62 | 1978.64 | 1805.88 |
| *D. mauritiana* | 10 | 1379.61 | 1798.25 | 1911.56 | 1696.47 |
| *D. santomea* | 10 | 1358.88 | 1693.15 | 1785.28 | 1612.44 |
| *D. yakuba* | 10 | 1368.38 | 1739.29 | 1805.78 | 1637.82 |
| *D. erecta* | 10 | 1352.29 | 1680.74 | 1820.83 | 1617.95 |
| *D. bipectinata* | 10 | 1326.81 | 1677.58 | 1712.40 | 1572.27 |
| *D. ananassae* | 10 | 1483.34 | 1865.13 | 1892.38 | 1746.95 |
| *D. persimilis* | 10 | 1601.28 | 1893.45 | 2083.31 | 1859.35 |
| *D. willistoni* | 10 | 1403.30 | 1737.83 | 1771.76 | 1637.63 |
| *D. paramelanica* | 10 | 1852.78 | 2267.74 | 2379.89 | 2166.80 |
| *D. melanica* | 10 | 1809.51 | 2266.98 | 2386.99 | 2154.49 |
| *D. mojavensis* | 10 | 1444.44 | 1678.11 | 1800.84 | 1641.13 |
| *D. buzzatii* | 10 | 1623.25 | 1952.49 | 2015.45 | 1863.73 |
| *D. hydei* | 10 | 2177.33 | 2671.00 | 2772.14 | 2540.16 |
| *D. novamexicana* | 10 | 1930.75 | 2380.17 | 2529.14 | 2280.02 |
| *D. americana* | 10 | 2042.64 | 2501.99 | 2657.44 | 2400.69 |
| *D. virilis* | 10 | 2172.87 | 2637.23 | 2910.11 | 2573.40 |
| *D. immigrans* | 10 | 2357.48 | 2790.73 | 2893.47 | 2680.56 |

**Table S4-Related to Figure 4: Mean body size of all 20 species for male flies.** Body sizes were calculated from tracked videos and averaged for each species.

| **Species** | **Sample Size** | **Mean Body Size (µm)** |
|---|---|---|
| *Ch. procnemis* | 20 | 758.85 |
| *D. melanogaster* | 46 | 625.95 |
| *D. sechellia* | 22 | 557.24 |
| *D. mauritiana* | 26 | 540.98 |
| *D. santomea* | 23 | 491.66 |
| *D. yakuba* | 24 | 502.41 |
| *D. erecta* | 25 | 489.90 |
| *D. bipectinata* | 22 | 465.98 |
| *D. ananassae* | 23 | 514.88 |
| *D. persimilis* | 25 | 634.13 |
| *D. willistoni* | 23 | 522.65 |
| *D. paramelanica* | 23 | 734.58 |
| *D. melanica* | 22 | 749.40 |
| *D. mojavensis* | 25 | 625.39 |
| *D. buzzatii* | 27 | 767.20 |
| *D. hydei* | 25 | 856.84 |
| *D. novamexicana* | 24 | 860.91 |
| *D. americana* | 27 | 889.59 |
| *D. virilis* | 25 | 866.36 |
| *D. immigrans* | 23 | 791.03 |

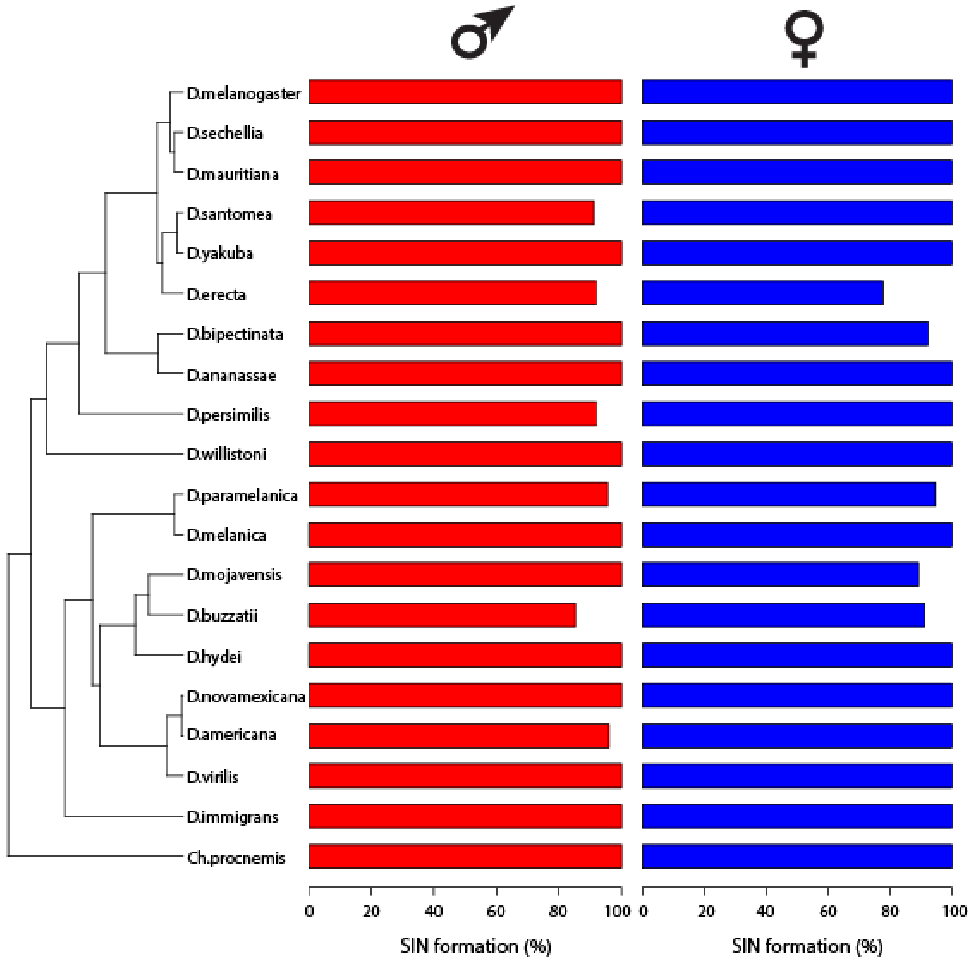


**Figure S1-Related to Method Details and Figure 3: Ability to form SINs for the male dataset (red) and female dataset (blue).** The x-axis is expressed as a percentage of the number of videos with at least one SIN iteration divided by the total number of videos acquired for each species. There are no differences across species in their ability to form networks ($p = 1$; $\chi 2$ goodness of fit test). The total number of videos that formed SINs are as follows: *D. melanogaster*: n= 46 (male), n=48 (female); *D.sechellia*: n=22 (male), n=22 (female); *D. mauritiana*: n=26 (male), n=27 (female); *D. santomea*: n=21 (male), n=23 (female); *D. yakuba*: n=24 (male), n=26 (female); *D.erecta*: n=23 (male), n=21 (female); *D. bipectinata*: n=22 (male), n=24 (female); *D. ananassae*: n=23 (male), n=25 (female); *D. persimilis*: n=23 (male), n=25 (female); *D. willistoni*: n=23 (male), n=24 (female); *D.paramelanica*: n=22 (male), n=18 (female); *D. melanica*: n=22 (male), n=21 (female); *D. mojavensis*: n=25 (male), n=25 (female); *D. buzzatii*: n=23 (male), n=21 (female); *D. hydei*: n=25 (male), n=24 (female); *D. novamexicana*: n=24 (male), n=22 (female); *D. virilis*: n=25 (male), n=27 (female); *D. immigrans*: n=23 (male), n=22 (female); *Ch. procnemis*: n=20 (male), n=21 (female).

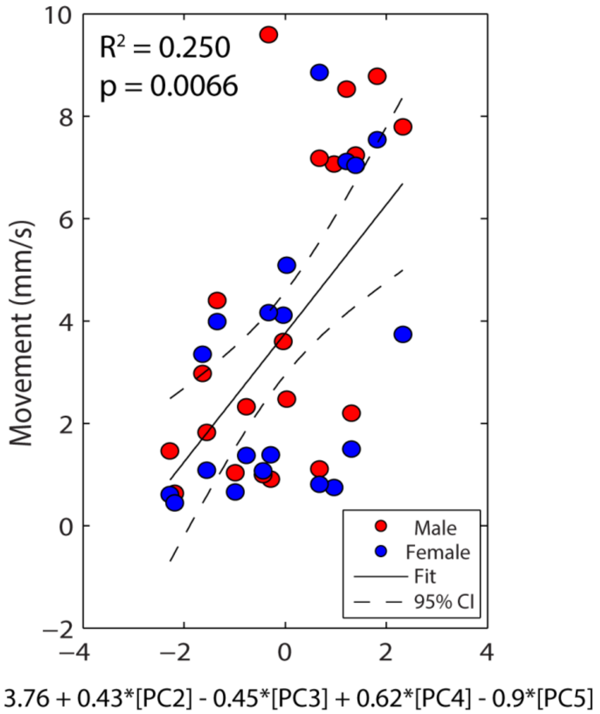


**Figure S2-Related to Figure 5: Environmental model for the level 1 variable Movement.** For the regression, each data point represents the mean SIN measure for a single species. The mean SIN measure for groups of male flies and female flies were pooled into each regression and are labeled with red and blue points, respectively. The solid trend line indicates line of best fit and dashed lines indicate 95% confidence interval of the model. The equation for this model is listed below the x-axis.

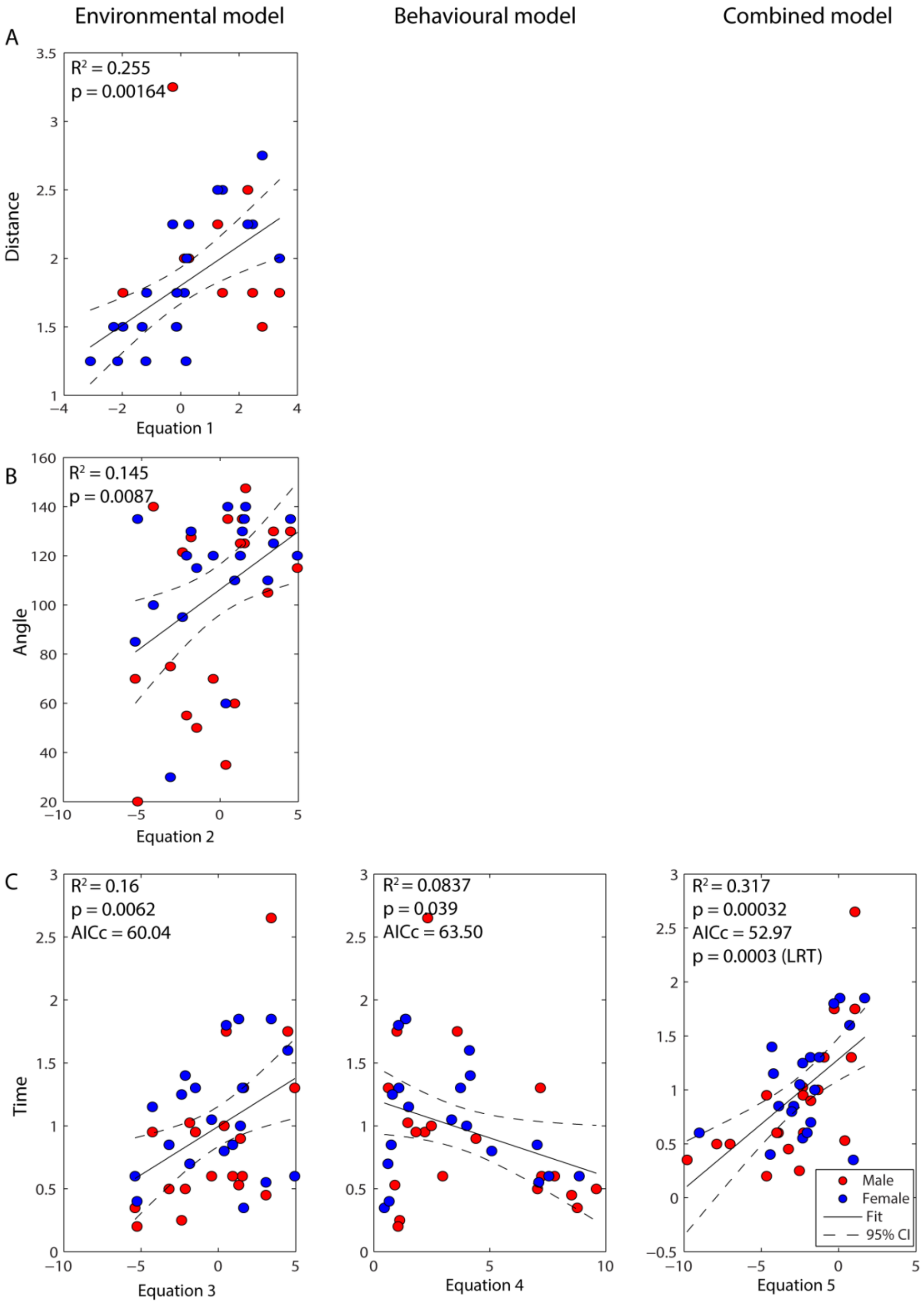


**Figure S3-Related to Figure 5: Environmental, behavioural and combined models for the level 2 social spacing variables: Distance, Angle and Time.** For all regressions, each data point represents the mean SIN measure for a single species. The mean SIN measure for groups of male flies and female flies were pooled into each regression and are labeled with red and blue points, respectively. Each solid trend line indicates line of best fit and dashed lines indicate 95% confidence interval of the model. **A)** No significant behavioural model formed for distance, therefore no combined model to report. The environmental model (Equation 1 predicts social distance ($R^2 = 0.255$, $p = 0.00164$). **B)** No significant behavioural model formed for angle, therefore no combined model to report. The environmental model (Equation 2) predicts angle ($R^2 = 0.145$, $p = 0.0087$). **C)** The environmental model (Equation 3) predicts time ($R^2 = 0.16$, $p = 0.0062$). The behavioural model (Equation 4) predicts time ($R^2 = 0.0837$, $p = 0.039$). The combined model significantly improves prediction of time compared to the behavioural model alone (Combined AICc < Behavioural AICc; $p = 0.0003$, likelihood ratio test). **Equation 1:** y = 1.80 – 0.071*[PC1] – 0.13*[PC5]. **Equation 2:** y = 106.3 + 4.71*[PC1]. **Equation 3:** y = 0.99 + 0.077*[PC1]. **Equation 4:** y = 1.21 – 0.061*[movement]. **Equation 5:** y = 1.28 + 0.091*[PC1] – 0.078*[movement].

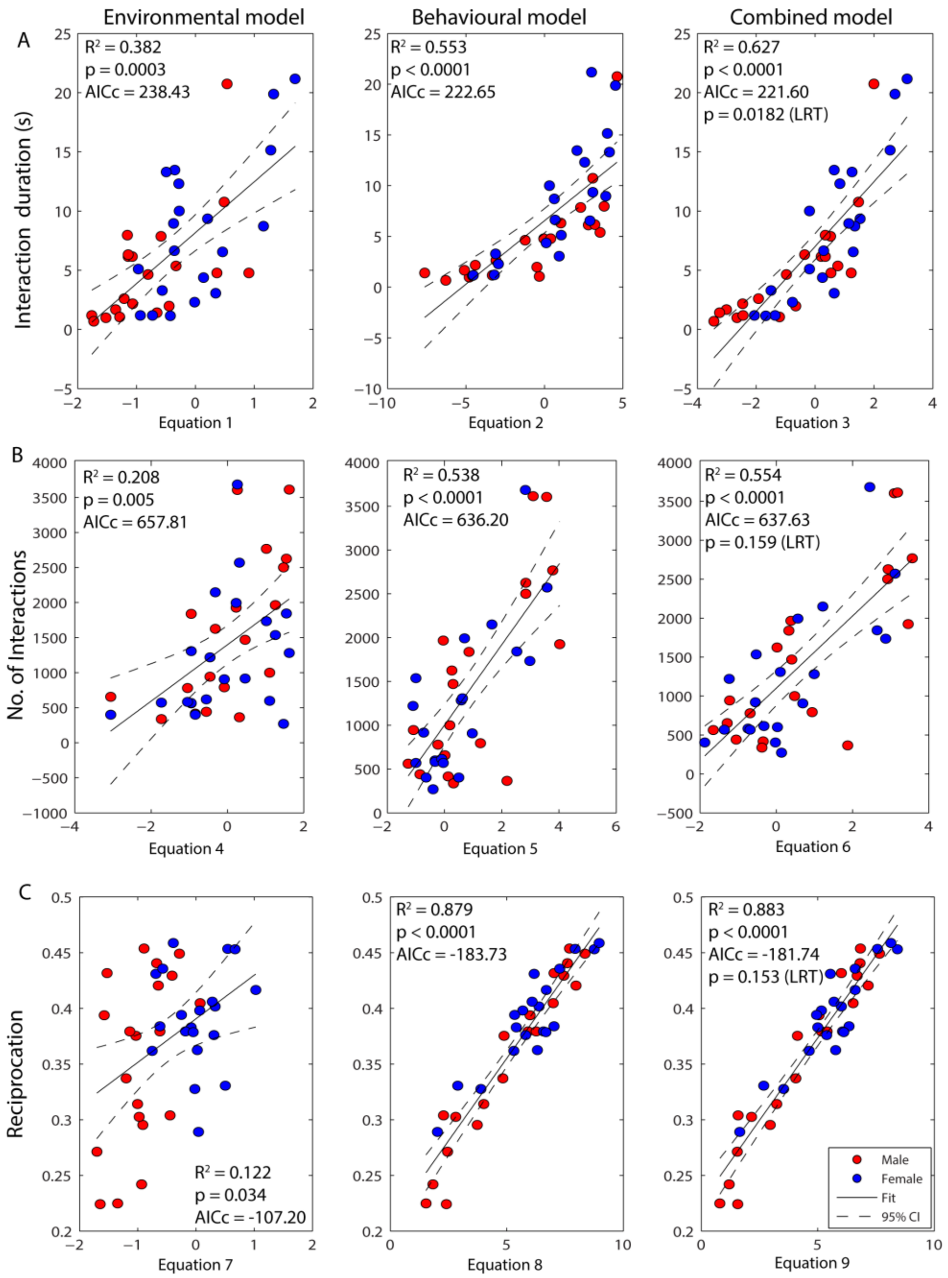


**Figure S4-Related to Figure 5: Environmental, behavioural and combined models for the level 3 variables interaction duration, number of interactions and reciprocation.** For all regressions, each data point represents the mean SIN measure for a single species. The mean SIN measure for groups of male flies and female flies were pooled into each regression and are labeled with red and blue points, respectively. Each solid trend line indicates line of best fit and dashed lines indicate 95% confidence interval of the model. **A)** The environmental model (Equation 1) predicts interaction duration ($R^2 = 0.382$, p = 0.0003). The behavioural model (Equation 2) predicts interaction duration ($R^2 = 0.553$, $p < 0.0001$). The combined model (Equation 3) does not significantly improve the prediction of interaction duration compared to the behavioural model alone (Combined AICc =~ Behavioural AICc; p = 0.0182, likelihood ratio test). **B)** The environmental model (Equation 4) predicts number of interactions ($R^2 = 0.20772$, p = 0.005). The behavioural model (Equation 5) predicts number of interactions ($R^2 = 0.538$, $p < 0.0001$). The combined model does not significantly improve the prediction of number of interactions compared to the behavioural model alone (Combined AICc =~ Behavioural AICc; p = 0.1594, likelihood ratio test). **C)** The environmental model (Equation 7) predicts reciprocation ($R^2 = 0.122$, p = 0.034). The behavioural model (Equation 8) predicts reciprocation ($R^2 = 0.879$, $p < 0.0001$). The combined model (Equation 9) does not significantly improve the prediction of reciprocation compared to the behavioural model alone (Combined AICc =~ Behavioural AICc; p = 0.153, likelihood ratio test). **Equation 1:** y = 8.17 – 0.76*[PC2] + 1.02*[PC3] + 2.31*[PC5]. **Equation 2:** y = 6.54 – 1.25*[movement] + 0.045*[angle]. **Equation 3:** y = 6.99 – 0.23*[PC2] + 0.47*[PC3] + 1.47*[PC5] – 0.93*[movement] + 0.038*[angle]. **Equation 4:** y = 1398 + 220*[PC4] – 336*[PC5]. **Equation 5:** y = 1001 + 213*[movement] – 406*[time]. **Equation 6:** y = 1100 + 91.5*[PC4] – 151*[PC5] + 183*[movement] – 387*[time]. **Equation 7:** y = 0.39 + 0.011*[PC3]. **Equation 8:** y = 0.21 – 0.0065*[movement] + 0.014*[social distance] + 0.0017*[angle] – 0.025*[time]. **Equation 9:** y = 0.22 + 0.0028*[PC3] – 0.0061*[movement] + 0.01*[social distance] + 0.0017*[angle] – 0.025*[time].

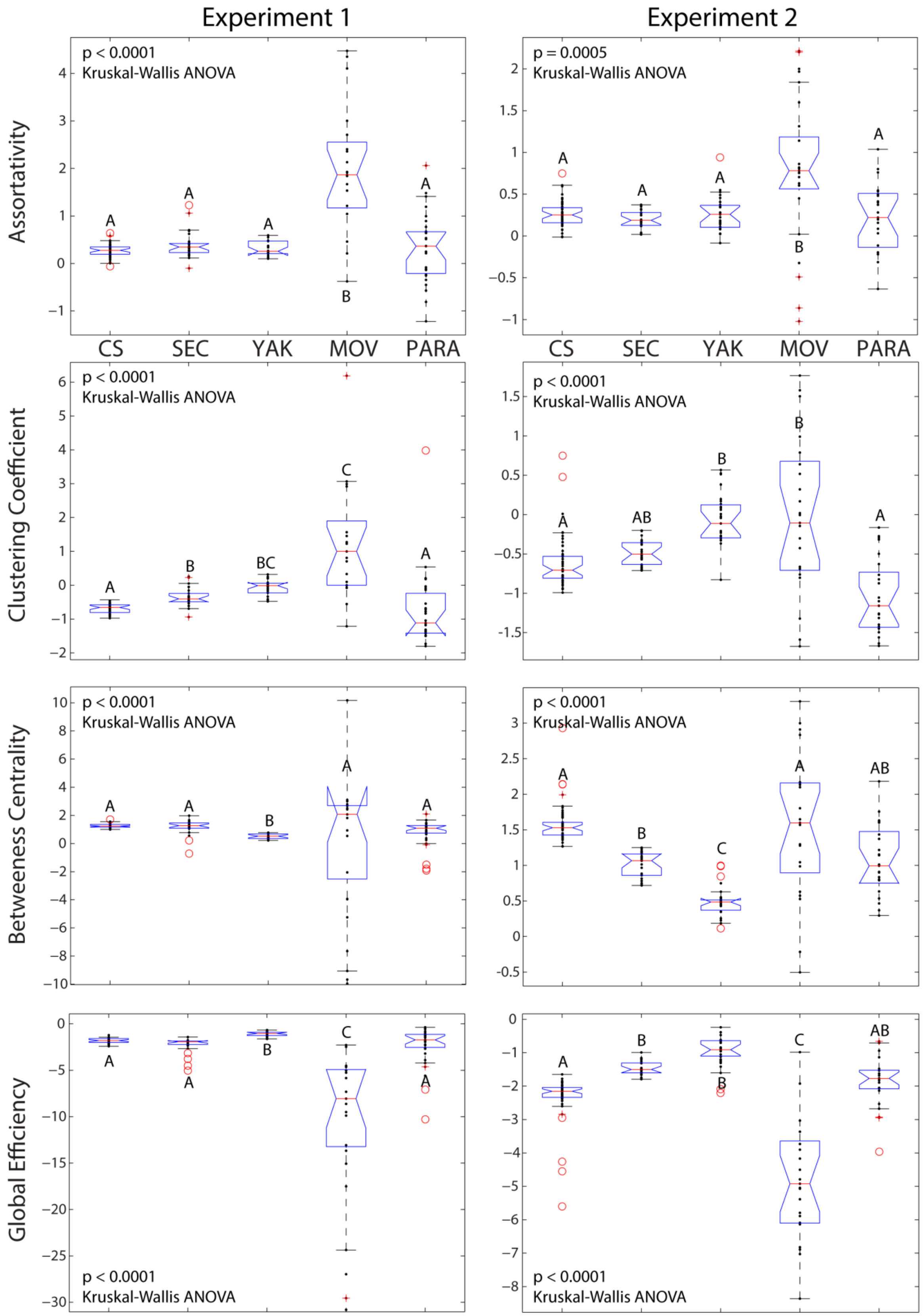


**Figure S5-Related to Figure 3: Relative species differences between *D. melanogaster* (CS), *D. sechellia* (SEC), *D. yakuba* (YAK), *D. mojavensis* (MOV), *D. paramelanica* (PARA) are replicated in male flies between pilot data (Experiment 1; left panels) and extended data (Experiment 2; right panels).** All figures are boxplots which outline the distribution of the z-scores for all four SIN measures. Letters above each box indicate statistically distinct groups from a Kruskal-Wallis one-way ANOVA followed Tukey-Kramer post-hoc tests.

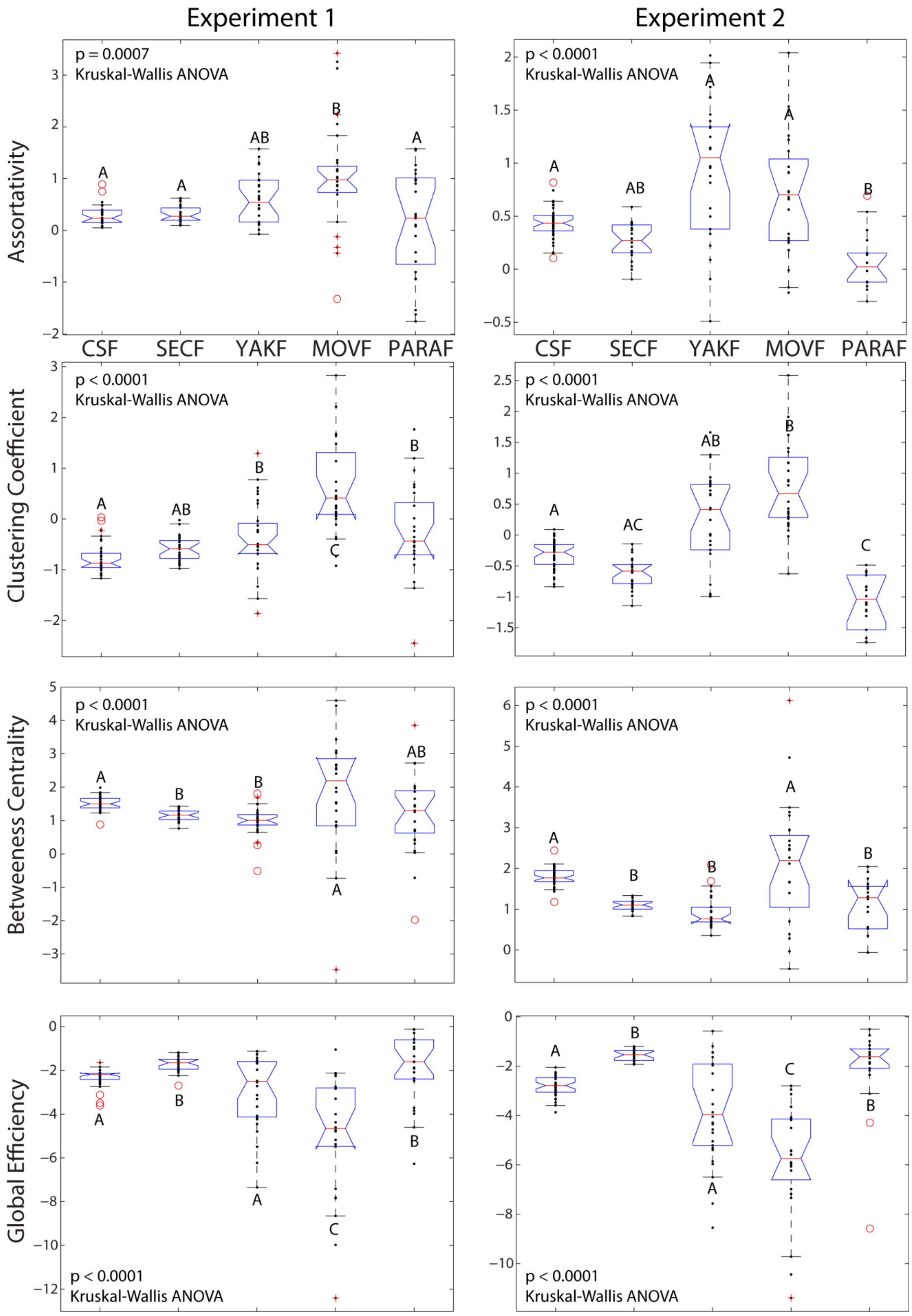


**Figure S6-Related to Figure 3: Relative species differences between *D. melanogaster* (CS), *D. sechellia* (SEC), *D. yakuba* (YAK), *D. mojavensis* (MOV), *D. paramelanica* (PARA) are replicated in female flies between pilot data (Experiment 1; left panels) and extended data (Experiment 2; right panels).** All figures are boxplots which outline the distribution of the z-scores for all four SIN measures. Letters above each box indicate statistically distinct groups from a Kruskal-Wallis one-way ANOVA followed Tukey-Kramer post-hoc tests.

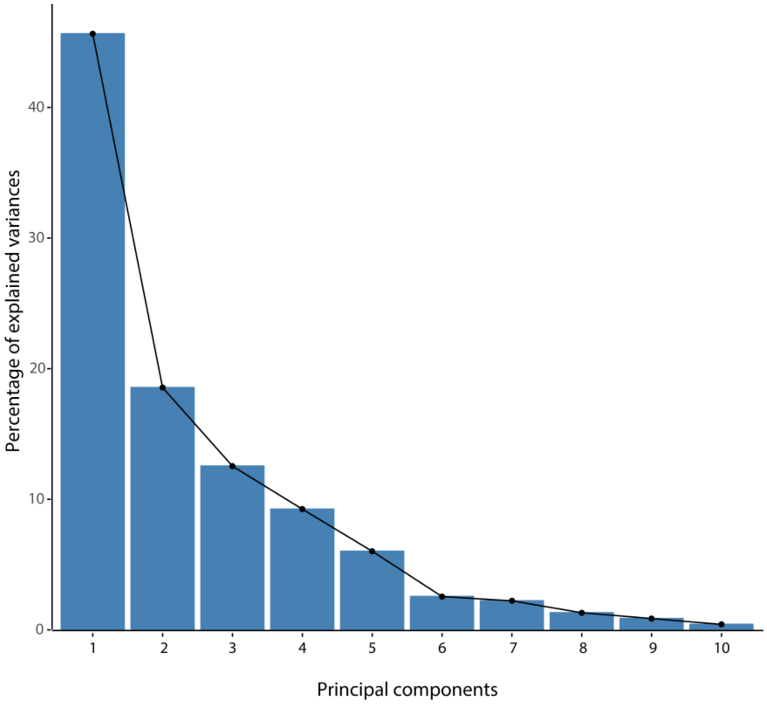


**Figure S7-Related to Figure 5: The first 5 principal components of the climatic measures, extracted from each species geographic origin, account for 92% of the variance across the 19 variables.** Each bar represents the percentage of variance explained by the first 10 dimensions. Because the rate of decrease reduces after dimension 5, the first 5 dimensions were used to represent the environmental variables for all regression analyses.
